# The Long Non-coding RNA Landscape of Endurance Exercise Training

**DOI:** 10.1101/2025.10.09.681231

**Authors:** Bernardo Bonilauri, Gregory R. Smith, Archana N. Raja, David Jimenez-Morales, Abdalla Ahmed, Christopher Jin, Lauren M. Sparks, Martin J. Walsh, Stephen B. Montgomery, the MoTrPAC Study Group, Sue C. Bodine, Euan A. Ashley, Maléne E. Lindholm

## Abstract

Long non-coding RNAs (lncRNAs) regulate multiple cellular processes. However, knowledge of the responses and regulatory functions of lncRNAs in physical exercise and training remains limited. As part of the Molecular Transducers of Physical Activity Consortium (MoTrPAC), we conducted a comprehensive analysis of lncRNA expression patterns in 18 tissues after an 8-week progressive endurance training program in rats. The lncRNA expression pattern was largely tissue-specific. In total, 759 unique lncRNAs were found to be differentially expressed across all tissues, generally displaying lower abundance, shorter transcript length, and reduced GC content compared with protein-coding genes. The most pronounced changes were observed in white and brown adipose tissues, the hypothalamus, and the adrenal gland. In the two skeletal muscle tissues investigated, only two lncRNAs were commonly differentially expressed. White and brown adipose tissues revealed a correlation between upregulated differentially expressed lncRNAs and coding genes associated with immune regulation. We identified substantial sex differences in the lncRNA regulatory landscape in response to exercise training. This comprehensive tissue-specific characterization of exercise-responsive lncRNAs opens new avenues for understanding exercise as molecular medicine and may inform the development of lncRNA-targeted therapeutics that harness the beneficial effects of exercise.

## INTRODUCTION

Regular physical exercise is recognized for its wide-ranging health benefits. Exercise affects nearly all organ systems, leading to physiological adaptations that positively influence health and reduce the risk of numerous diseases^1^. In recent years, several studies using multi-omics techniques have paved the way for a greater understanding of the molecular mechanisms involved in the adaptations induced by exercise. This regulation occurs at the DNA, RNA, and protein levels^2–5^. Consequently, “omics” techniques such as epigenomics, transcriptomics, and proteomics have proven invaluable in identifying changes in signaling pathways and cellular dynamics in response to different stimuli. For transcriptomics, recent meta-analyses have examined gene expression changes associated with physical activity and identify genes and pathways influenced by both exercise and inactivity^6^. Using linear mixed-effects meta-regression on 43 publicly accessible datasets comprising skeletal muscle and blood samples from individuals who underwent endurance and resistance exercise, Amar et al. demonstrated that transcriptional responses differ between acute exercise and training and vary when critical factors such as age and sex are taken into account^7^. Furthermore, several studies have showcased substantial alterations in the proteome and secretome of various tissues in response to exercise, thereby impacting multiple pathways associated with tissue physiology and metabolism^8–11^. However, previous studies are limited to mRNAs, thus there is still limited understanding of the expression and functions of non-coding RNAs (e.g., microRNAs, long non-coding RNAs, short RNAs, circular RNAs) during exercise and training.

Long non-coding RNAs (lncRNAs) are commonly defined as a class of non-coding transcripts longer than 200 nucleotides, comprising a substantial portion of the non-coding transcriptome. They exhibit diversity in terms of their size, function and origin, and their classification has been a subject of ongoing research and debate^12^. Many lncRNAs undergo splicing and polyadenylation, resembling messenger RNAs (mRNAs), while others lack polyadenylation and 7-methylguanosine capping. These RNAs play diverse regulatory and architectural roles in the cytoplasm and nucleus, often exhibiting tissue-specific expression patterns and limited conservation across species^12,13^. LncRNAs are involved in epigenetic and transcriptional regulation, splicing, and translational control, and regulate pathways essential for cell development, growth, differentiation, and proliferation^12–15^. They have also demonstrated utility as biomarkers for several conditions and diseases, highlighting their potential in biomedical research^16–18^. In the context of skeletal muscle, specific lncRNAs have been implicated in myogenesis, myoblast proliferation, and muscle performance^19^. More recently, studies have revealed differential expression of lncRNAs in skeletal muscle response to exercise training, including in humans, indicating their potential involvement in exercise-induced adaptations^20–22^. With the goal of developing a molecular map of the response to exercise and training, the Molecular Transducers of Physical Activity Consortium (MoTrPAC)^1^ recently published a comprehensive study of the temporal multi-omics response of adult male and female rats undergoing up to 8 weeks of incremental endurance training. Multi-tissue molecular responses to exercise were identified at different levels of regulation, including transcriptional, post-transcriptional, translational and metabolic, important in immunological, metabolism and stress response pathways^23^. While the original MoTrPAC study focused on protein-coding genes, lncRNAs were not systematically analyzed. Here, we conducted a de novo computation analysis of the transcriptome data, implementing realignment to the updated genome annotation and lncRNA identification pipelines to characterize the previously unexplored lncRNA landscape. Our analyses reveal tissue-specific differential expression patterns, with greater changes in adipose tissues, hypothalamus and adrenal glands. The differential lncRNAs are structurally different from mRNAs and we identify sex differences in the lncRNA regulatory landscape in response to exercise training, with implications for adaptation mechanisms and the potential for utilizing the non-coding genome for therapeutic development.

## RESULTS

### Endurance Exercise Training Triggers Tissue-and Sex-Specific lncRNA Reprogramming

To systematically map training-responsive lncRNAs, we utilized RNA sequencing (RNA-seq) data from a previously published comprehensive study on the temporal, multi-omics responses to endurance exercise training across 18 tissues in adult male and female rats^23^. These rats either remained sedentary or underwent progressive exercise training for 1, 2, 4, and 8 weeks. After 8 weeks, both male and female rats exhibited a significant increase in maximal aerobic capacity (VO₂max). Male rats showed a marked reduction in body fat percentage, whereas female rats did not^23^.

The availability of multi-omics data from these rats provided an opportunity to study lncRNA responses in exercise training across a range of tissues, which remain largely unexplored. To address this gap, we performed a comprehensive omics analysis using datasets from 18 tissues: adrenal, brown adipose tissue (BAT), blood, colon, cortex, heart, hippocampus, hypothalamus, kidney, liver, lung, ovary, *gastrocnemius* muscle (SKMGN), *vastus lateralis* muscle (SKMVL), small intestine (SMLINT), spleen, testes, and white adipose tissue (WAT) (**Figure 1A**). To better capture lncRNA expression changes induced by endurance training, we first reprocessed the RNA-seq data using the updated rat genome assembly (mRatBN7.2) and subsequently focused on identified lncRNAs that exhibit an FPKM (fragments per kilobase of transcript per million mapped reads) value > 1 (see Materials and Methods). **Figure 1B** illustrates the count of uniquely expressed lncRNAs over the course of the 8-week training period, demonstrating the highest lncRNA expression in testes, confirming previous findings^24,25^, and the lowest in whole blood. Differential expression analysis of lncRNAs (DELncs; using a significance threshold of *p* < 0.01 and log2 fold-change |log2FC| > 1) identified 759 unique differential lncRNAs across all tissues (**Figure 1C**). White adipose tissue (87 DELncs), hypothalamus (86 DELncs), brown adipose tissue (73 DELncs), and adrenal glands (65 DELncs) exhibited the most pronounced responses, while testis showed minimal changes (6 DELncs). Temporal patterns varied dramatically between tissues with some peaking early (small intestine at week 1), while others showed delayed responses (adrenal gland and blood at week 8), suggesting tissue-autonomous regulatory programs rather than systemic coordination (**Figure 1D**).

**Figure 1.**
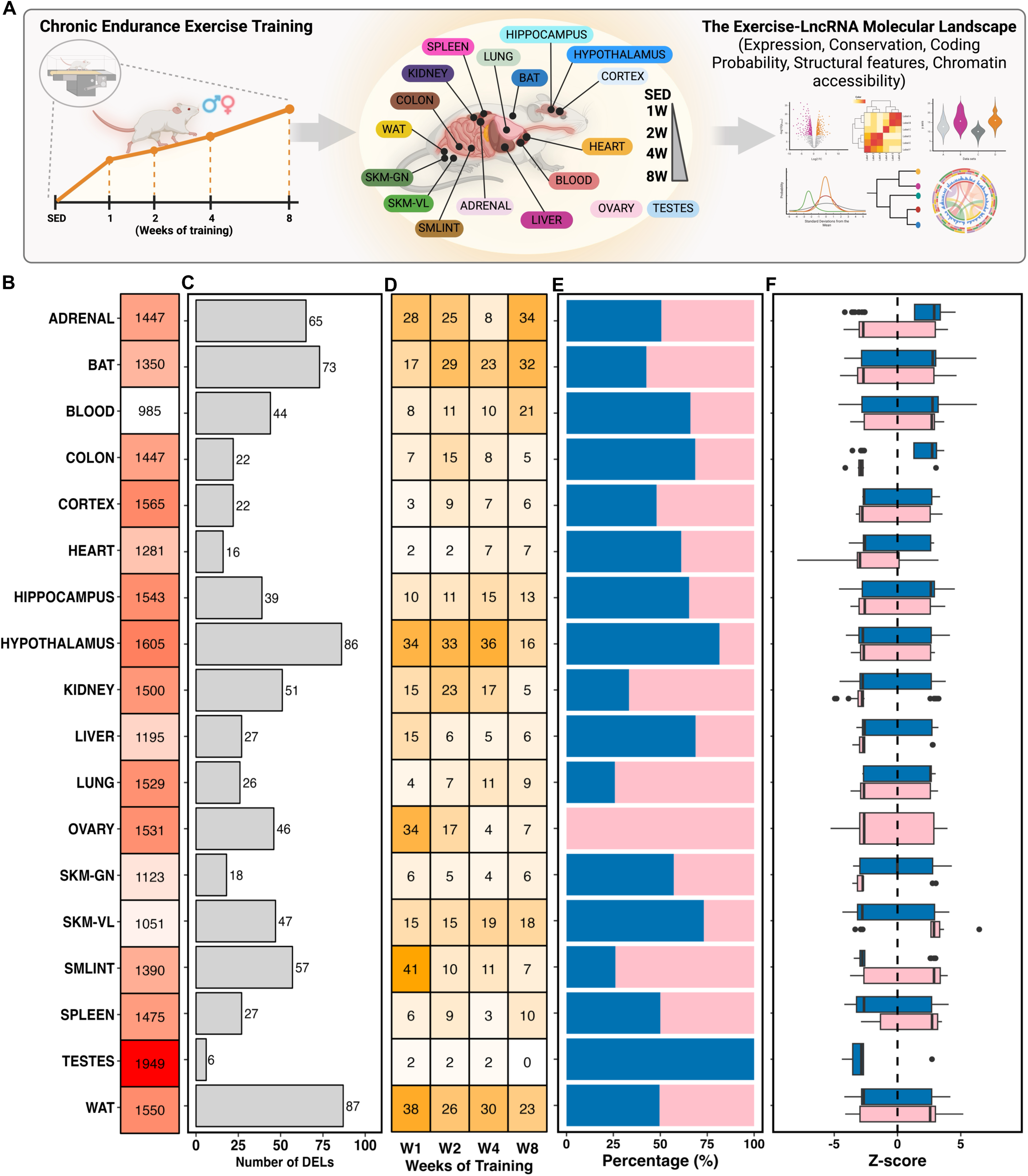
The landscape of long non-coding RNAs (lncRNAs) in endurance exercise training. **(A)** Schematic overview of the MoTrPAC-lncRNA project. Male and female 6-month-old rats were subjected to progressive endurance exercise training (i.e., treadmill), with 18 tissues collected from sedentary animals or animals completing 1, 2, 4, and 8 weeks of training. Tissues were processed and subjected to multi-omics techniques and analysis. The lncRNAs were analyzed at transcriptional and epigenomic levels. **(B)** The first panel shows the total number of unique identified lncRNAs (FPKM > 1) in each tissue after 8 weeks of aerobic training. **(C)** Among the highlighted lncRNAs, we identified differentially expressed lncRNAs (DELncs, |log2FC| >1, *p* < 0.01). **(D)** The DELncs exhibited a distinct expression pattern for each analyzed tissue, as well as across different weeks of training. **(E-F)** The distribution of DELncs by sex (male in blue, female in pink) and the average expression in each tissue were shown, respectively.

Interestingly, sex-specific differences dominated the differential lncRNA landscape. Only seven tissues exhibited differential lncRNAs common to both sexes, accounting for just 21% of lncRNAs in the adrenal gland, 5% in BAT, 9% in the colon, 6% in the heart, 1% in the hypothalamus, 2% in the kidney, and 4% in *vastus lateralis* (**Figure S1A**). Certain tissues show a similar number of differential lncRNAs between males and females (e.g., WAT, spleen, cortex, and adrenal), while others display a higher prevalence of sex-specific differential lncRNAs (e.g., hypothalamus, small intestine, liver, lung, and kidney) (**Figure 1E**). Even when lncRNAs were differentially expressed in both sexes, they frequently showed opposing patterns or occurred at different time points (**Figure 1F, Figure S1B**), underscoring the complexity of sex-specific molecular responses to exercise. Blood, cortex, hippocampus, liver, lung, *gastrocnemius* and WAT, for example, displayed sex-exclusive differential lncRNA expression in response to training (**Figure S2**). The implications of these findings are particularly significant, as previous studies have documented that certain lncRNAs perform distinct functions and operate through different mechanisms in males and females^26–28^. Moreover, a recent study has shown that lncRNAs are significantly more likely to exhibit sex-biased expression than protein-coding genes in human skeletal muscle^29^. Notably, most multi-omics studies in animal models use a single sex for greater homogeneity, often neglecting sex-specific variability in lncRNA gene regulation.

### Tissue-Specific Clustering and Limited Evolutionary Conservation Define Differential lncRNA Landscape

To uncover non-linear patterns within the dataset, we leveraged t-distributed Stochastic Neighbor Embedding (t-SNE) and observed predominantly tissue-specific clustering with enhanced separation after 8 weeks of training in tissues such as BAT, spleen, lung, and *gastrocnemius* in males, and tissues such as colon, WAT, hypothalamus, and spleen in females (**Figure 2A and 2B**). To complement these observations with a linear approach, Principal Component Analysis (PCA) confirmed tissue-specificity as the largest source of variance, with secondary variation attributable to training duration (i.e., weeks of training) (**Figure S3A and S3B**). This pattern persisted across all differential lncRNAs and became more evident when eight internal organs (adrenal, colon, heart, kidney, liver, lung, small intestine, and spleen) exclusively (**Figure S3C and S3D**), aligning with the regulatory role of lncRNAs in differentiating tissue function.

**Figure 2.**
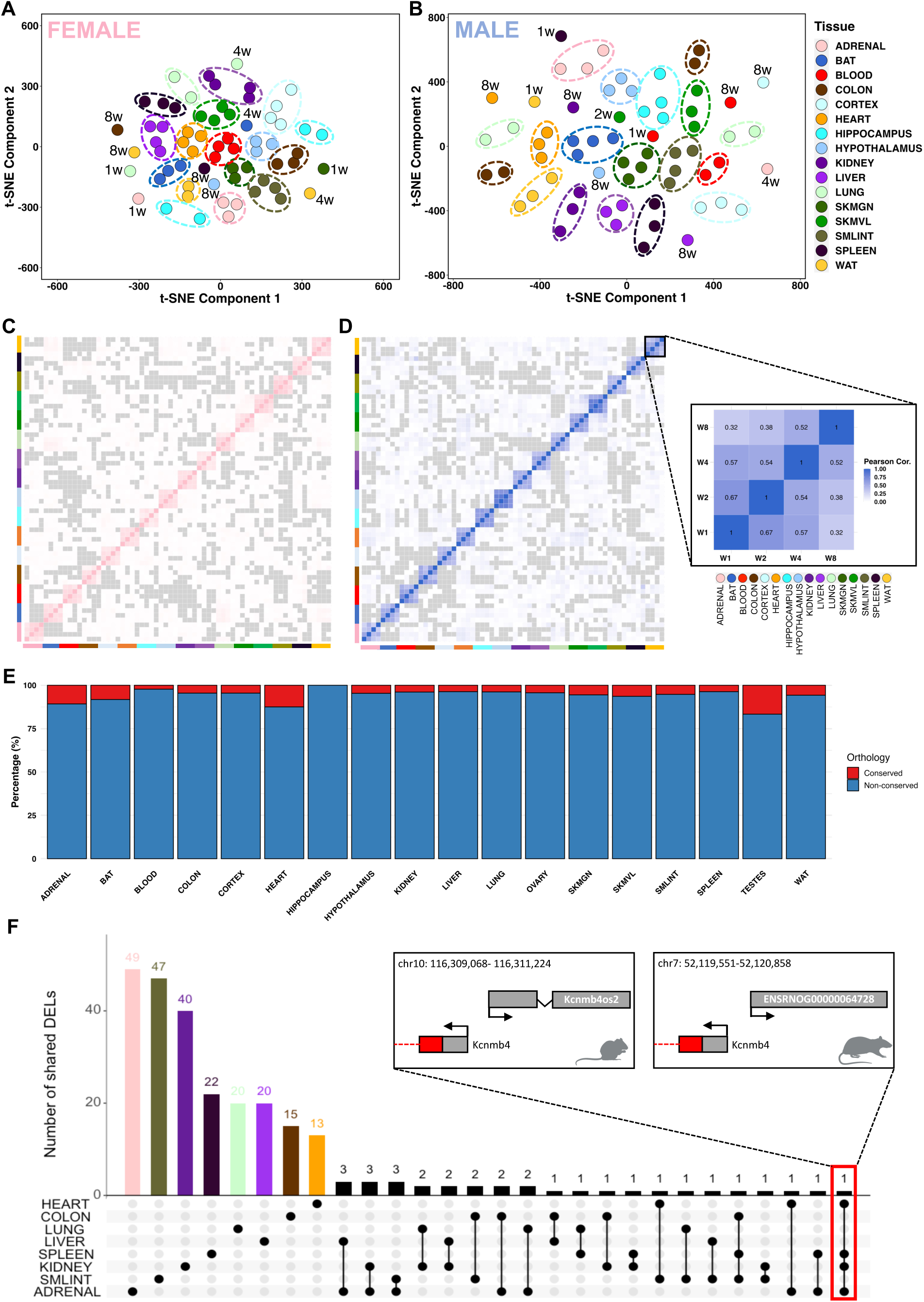
Tissue-specific lncRNA response to endurance training. **(A-B)** t-SNE analysis of all identified lncRNAs reveals clustering driven primarily by tissue type, with distinct patterns observed in female and male rats, respectively. **(C-D)** Pearson correlation heatmaps showing tissue-specific lncRNA expression across all training weeks in female (pink) and male (blue) rats, respectively. **(E)** Percentage of conserved and non-conserved differentially expressed lncRNAs (DELncs) orthologs between rat and human, identified using the Rat Genome Database (RGD), across all tissues. **(F)** The limited overlap of shared DELncs among the eight internal organs highlights the tissue-specific regulation of lncRNAs in response to endurance training, with genomic context provided for mouse-rat conserved DELncs such as Kcnmb4os2 and ENSRNOG00000064728.

To further explore the timewise lncRNA dynamics in response to endurance training, we next examined the correlation between the expression of all lncRNAs across the four training timepoints in all tissues, separated by sex (**Figure 2C and 2D**). We also analyzed the correlation for all expressed lncRNAs and differential lncRNAs independently (**Figure S3E and S3F**, respectively). There was a strong intra-tissue correlation across training weeks, reflecting remarkable consistency within individual tissues, and minimal inter-tissue correlation, underscoring the tissue-specific expression profiles. Analysis of lncRNA chromosomal origin showed no significant patterns (**Figure S4**).

Although conservation often indicates functionality, the absence of sequence conservation does not necessarily imply the opposite. This distinction becomes evident when comparing lncRNAs with protein-coding genes, as lncRNAs often exhibit distinct patterns of evolutionary conservation, which frequently involve lower levels of conservation between species^30,31^. Using the Rat Genome Database (RGD), we curated all differential lncRNAs, focusing on those that share orthologs with human genes. We found that the majority do not have orthologs between rats and humans (**Figure 2E**). The adrenal, heart, and testes displayed a higher proportion of conserved differential lncRNAs, while blood and hippocampus had minimal conservation. Notable conserved transcripts included the well-studied lncRNA Fendrr (ENSRNOG00000062673), down-regulated in kidney after 4 weeks of training, known to be essential for perinatal survival and predicted to have core promoter sequence-specific DNA binding activity^32^. The lncRNA ENSRNOG00000064053, previously known as LINC01314 or LOC102553298, was recently reannotated as protein_coding (CTXND1) through inference in humans due to its encoding capacity of a 59-amino acid protein. Another conserved transcript identified as the Ufm1 gene (ENSRNOG00000038176; Ensembl/RGD: lncRNA/protein_coding), predicted to be involved in various cellular processes, including endoplasmic reticulum function and cellular stress response, was found to be downregulated in heart, blood, ovary, and testes. Furthermore, the adipose-related lncRNA ENSRNOG00000016480, identified as the *Acbd7* gene (Ensembl/RGD: lncRNA/protein_coding), plays a role in fatty acid metabolic processes and facilitates fatty-acyl-CoA binding activity, and was observed here to be upregulated in WAT. The lncRNA ENSRNOG00000016535 (Ensembl/RGD: lncRNA/protein_coding), which has been identified as the protein-coding gene *Ccl22*, exhibited down-regulation in both BAT and WAT following training. CCL22 is a cytokine involved in immunoregulatory and inflammatory processes. Notably, its levels are significantly elevated in the circulation of morbidly obese patients^33^. This high expression is positively correlated with BMI. Furthermore, increased expression of CCL22 has been observed in visceral adipose tissue when compared to subcutaneous adipose tissue of morbidly obese patients^33^. The predicted conservation varies at both the RNA and protein levels. For instance, rat *Ccl22* canonical open reading frame (ORF) shows 89% and 65% conservation with the protein coding ORF of mice and humans, respectively (**Figure S5A**). For the predicted *Acbd7* transcript, the rat canonical ORF and RNA transcript are not conserved when compared to both mice and humans (**Figure S5B**), while *Atp5mc1* reaches 98% mouse and 87% human sequence conservation (**Figure S5C**).

When examining differential lncRNAs for the eight internal tissues exclusively, we confirmed the tissue-specific expression patterns, with only one lncRNA identified as common to four of the analyzed tissues (**Figure 2F**). The shared lncRNA, found in adrenal, kidney, spleen, and heart, is located antisense to the protein-coding gene *Kcnmb4* and lacks human conservation but shows homology with mouse lncRNA *Kcnmb4os2* (**Figure 2F; Figure S7A**). One of the regulatory roles of lncRNAs is to function as competing endogenous RNA (ceRNA), sequestering microRNAs (miRNAs). We identified 10 miRNAs that bind to the homologous region of *Kcnmb4os2* in mice (miR-217, miR-338-3p, miR-221-3p, miR-222-3p, miR-24-3p, miR-375-3p, miR-183-5p, miR-103-3p, miR-107-3p, miR-455-3p). Next, we retrieved all the target mRNAs of these miRNAs in rats from the miRDB database and cross-referenced them with the differentially expressed upregulated protein-coding genes in the four target tissues and the corresponding training week of our dataset (see Materials and Methods). This process resulted in the construction of a lncRNA-miRNA-mRNA network (**Figure S7**), revealing tissue-specific pathway enrichments: a cardiac network involved in positive regulation of cellular catabolism and autophagy processes (**Figure S7B**), a splenic network regulating protein and cellular metabolism (**Figure S7C**), and an adrenal network, which only showed general molecular functions (**Figure S7D**).

### Skeletal Muscle Subtypes Deploy Distinct lncRNA Regulatory Programs

Skeletal muscle is a plastic tissue that undergoes extensive molecular and structural adaptations to exercise training^34–37^. Despite their shared function in locomotion, skeletal muscles differ markedly in their responses to exercise, influenced by fiber composition, morphology, activation, and training mode^38,39^. Here, *vastus lateralis* and *gastrocnemius* muscles demonstrated remarkable lncRNA transcriptional variability, primarily explained by the duration of the training regimen (i.e., weeks of training) and sex (**Figure 3A; Figure S8A and S8B**). Average lncRNA expression increased over the eight weeks of training in *vastus lateralis* (males: mean Z-score= -1.05, ι1Z-score(w8–w1)= +2.85; females: mean Z-score= 1.91, ι1Z-score(w8–w1)= +3.29). In *gastrocnemius,* males showed a reduction (mean Z-score= 0.001, ι1Z-score(w8–w1)= -3.36), whereas females, despite a positive change from week 1 to week 8 (ι1Z-score(w8–w1)= +2.94), still exhibited an overall negative mean expression (mean Z-score= -1.68) (**Figure 3B**). When comparing the sets of differentially expressed lncRNAs, only two transcripts were shared between the two muscles (**Figure 3C**); lncRNA ENSRNOG00000069758 was downregulated in *gastrocnemius* after 1 week but upregulated in *vastus lateralis* after 8 weeks of training, whereas lncRNA ENSRNOG00000063951 was downregulated in female *gastrocnemius* at 4 weeks and in male *vastus lateralis* after 8 weeks (**Figure S8C**). Consequently, 96% and 89% of the differential lncRNAs in *vastus lateralis* and *gastrocnemius*, respectively, exhibit muscle subtype specificity.

**Figure 3.**
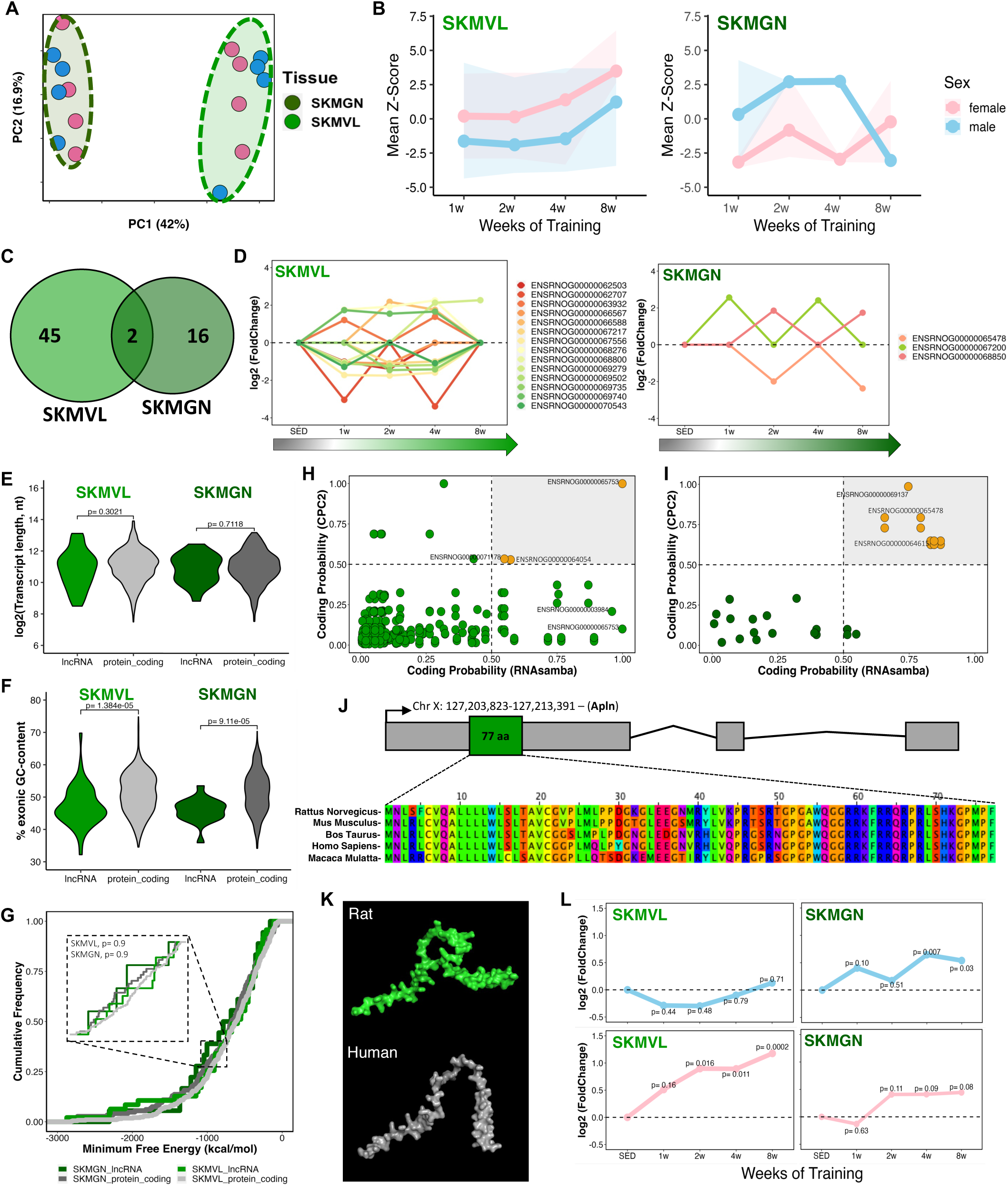
The transcriptional landscape of lncRNAs in skeletal muscle after 8 weeks of aerobic exercise. **(A)** Principal Component Analysis (PCA) demonstrating the clustering pattern of all identified lncRNAs in the gastrocnemius (SKMGN, dark green) and vastus lateralis (SKMVL, light green) across different training weeks. **(B)** Mean Z-score distribution of all differentially expressed lncRNAs (DELncs) in SKMVL and SKMGN during the 8-week training regimen, stratified by sex (male rats in blue, female rats in pink). **(C)** Venn diagram illustrating the overlap of DELncs between SKMVL and SKMGN, with most DELncs being unique to each tissue. **(D)** Gene expression changes (log2FC) of DELncs identified across multiple training points in SKMVL and SKMGN, respectively. **(E)** Comparison of the length in nucleotides (nt) of DELncs and differentially expressed protein_coding genes (DEGs) in SKMVL (light green) and SKMGN (dark green). **(F)** Comparison of the exonic GC content percentage in SKMVL (light green) and SKMGN (dark green). **(G)** Analysis of Minimum Free Energy (MFE) of identified DELncs and DEGs in each tissue revealed no significant statistical difference. **(H-I)** Scatter plots illustrating the analysis of protein-coding potential of DELncs in SKMVL and SKMGN, respectively. **(J)** Identification of the DELnc ENSRNOG00000003984 as the Apelin gene, partially conserved across species. **(K)** 3D structural representation of the Apelin protein in rat (green) and human (gray). The rat model (UniProt Q9R0R3, isoform 1) and the human model (UniProt Q9ULZ1, isoform 1) correspond to AlphaFold v4 predicted structures, retrieved from the AlphaFold Protein Structure Database (DeepMind and EMBL-EBI). **(L)** Apelin expression is observed to increase after 8 weeks of endurance training. Blue line represents male rats and pink line represent female rats.

Among all differential lncRNAs, those detected at multiple training time points may represent more robust and consistent transcriptional changes. This was observed for 14 lncRNAs in *vastus lateralis* and only 3 transcripts for *gastrocnemius* (**Figure 3D**), suggesting a regulatory role across the training time course that can be related to skeletal muscle adaptation to endurance exercise. Another key aspect of lncRNA biology is their structural comparison to protein-coding genes. Typically, lncRNAs exhibit smaller transcript sizes than mRNAs, lower GC content, higher minimum free energy, and overall lower expression levels, reflecting reduced structural constraint, sequence conservation, and abundance relative to protein-coding genes, often in a more cell type-or tissue-specific manner^12,40,41^. Here, in muscles, the differential lncRNAs did not differ in length compared to mRNAs (**Figure 3E**) but had significantly lower GC content (**Figure 3F**) and minimum free energy (MFE) values similar to those of differential mRNAs (DEGs) (**Figure 3G**). Furthermore, these lncRNAs were expressed at significantly lower levels than protein-coding genes in *gastrocnemius*, but not in *vastus lateralis*. (**Figure S9M and S9N**). These comparative analyses were performed for all tissues (**Figure S9 and S10**). The absence of a distinct compositional pattern among these lncRNAs in rats highlights the complexity of predicting their potential functions and mechanisms of action, limiting the understanding of their biological role.

To unveil potential pathways under the regulation of these lncRNAs throughout the training weeks, we performed a correlation analysis between differential lncRNAs and mRNAs (DEGs) (see Materials and Methods). We identified enrichments solely for the correlated downregulated mRNAs in *gastrocnemius*, revealing pathways such as negative regulation of myoblast differentiation, myelination, anatomical structure development, skeletal muscle cell differentiation, and developmental process (**Figure S8D**). These pathways are intricately linked to the physiological adaptations of skeletal muscles to endurance training.

Next, we undertook analyses of protein-coding capability based on the sequence of differential RNA transcripts and all isoforms, with 94% and 83% identified as non-coding in *vastus lateralis* and *gastrocnemius*, respectively (**Figure 3H and 3I**). LncRNAs with a score greater than 0.5 were classified as having protein-coding potential^42,43^, possibly being misannotated or harboring small open reading frames (smORFs) capable of encoding microproteins^44^. This analysis was performed for all studied tissues and results are shown in **Figure S11**. Moreover, we performed a comprehensive scan and integration of data from these lncRNA sequences and our previously published proteomic data^23^ to identify potentially misannotated lncRNAs or microproteins encoded by smORFs. Only a small number of candidate peptides were detected, most of which were classified as false positives or reannotated – for instance, ENSRNOG00000067060, which has recently been updated to a protein-coding gene encoding an 83-aa protein (Nmes1; Uniprot Q5RK28).

Among the differential lncRNAs, we identified the lncRNA ENSRNOG00000003984 (Ensembl/RGD: lncRNA/protein_coding) as the partially conserved rat Apelin gene (APLN) that encodes for a 77-amino acid protein (**Figure 3J and 3K**). Apelin expression increased at several timepoints in *vastus lateralis* and *gastrocnemius* of both males and females (**Figure 3L**). To investigate if Apelin is also induced in human skeletal muscle with training, we analyzed public data from our previous meta-analysis comprising 1,725 samples collected from 739 individuals (https://www.extrameta.org)^7^. Apelin expression similarly increased in human skeletal muscle in individuals undergoing acute and chronic exercise (**Figure S8E and S8F**), and has been shown to be induced in myofibers in response to exercise in both mice and humans^45,46^. Aged mice that received daily apelin injections or underwent adenovirus-mediated gene transfer to overexpress apelin in skeletal muscle displayed enhanced muscle capacities and myofiber hypertrophy^45^. Apelin is also recognized as an important exerkine that triggers favorable metabolic effects in muscles, adipose tissue, and in various other target organs. For example, Apelin induces browning of white adipose tissue by increasing thermogenesis and promoting metabolic health, which is important for prevention of obesity and metabolic syndrome^47,48^. In WAT, Apelin increased only in the female animals following 8 weeks of endurance training (**Figure S12D**), while opposite changes were found in BAT. Dynamic Apelin expression could potentially contribute to adipose tissue remodeling and enhancements in its metabolic profile. Thus, we shed light on this critical transcript, identifying it as a protein-coding gene (i.e., Apelin) that might have been overlooked in previous rat transcriptomic studies due to its initial annotation as a long non-coding gene.

### Training-Regulated LncRNAs in Adipose Tissue Depots are Associated with Immune Modulation

In recent years, adipose tissue has emerged as a tissue profoundly affected by exercise training^49^. Different adipose tissue types exhibit intricate and distinct adaptations in response to exercise stimuli. Both brown adipose tissue (BAT), recognized for its thermogenic traits and energy dissipation capability, and white adipose tissue (WAT), primarily accountable for energy storage, undergo remodeling and metabolic alterations following exercise^50^. Both depots, that stem from different mesenchymal origins, function as endocrine organs, whose divergent responses contribute to the whole-body metabolic benefits of exercise. Similar to muscle tissues, PCA of the identified unique lncRNAs in BAT and WAT revealed tissue-specific clustering, with variations attributed to training durations (i.e., weeks of training) and sex (**Figure 4A, Figure S12A and S12B**). Differential lncRNA expression differed markedly between depots, where BAT showed increased average expression over the eight weeks of exercise in males (mean Z-score= 1.05, ι1Z-score(w8–w1)= +3.35), with a similar pattern observed in females, except for a decrease during the fourth week of training (mean Z-score= -0.57, ι1Z-score(w8–w1)= +4.69). In WAT, responses were sex-divergent: males exhibited lower overall levels and a net decrease (mean Z-score levels of -0.76, ι1Z-score(w8–w1)= -1.04), whereas females showed slightly higher overall levels and a net increase (mean Z-score 0.14, ι1Z-score(w8–w1)= +1.27) (**Figure 4B**). Only 5% of differential lncRNAs overlapped between depots (**Figure 4C, Figure S12C**), resulting in 89% (BAT) and 91% (WAT) showing tissue-specificity. Sustained regulation across multiple training time points occurred for 19 transcripts in BAT and 22 transcripts in WAT (**Figure 4D**), suggesting a potential regulatory role of these lncRNAs in adipose tissue adaptation to endurance training. This subset encompasses lncRNAs that exhibit highly significant regulation in BAT (i.e., ENSRNOG00000069911, ENSRNOG00000065889, and ENSRNOG00000064424).

**Figure 4.**
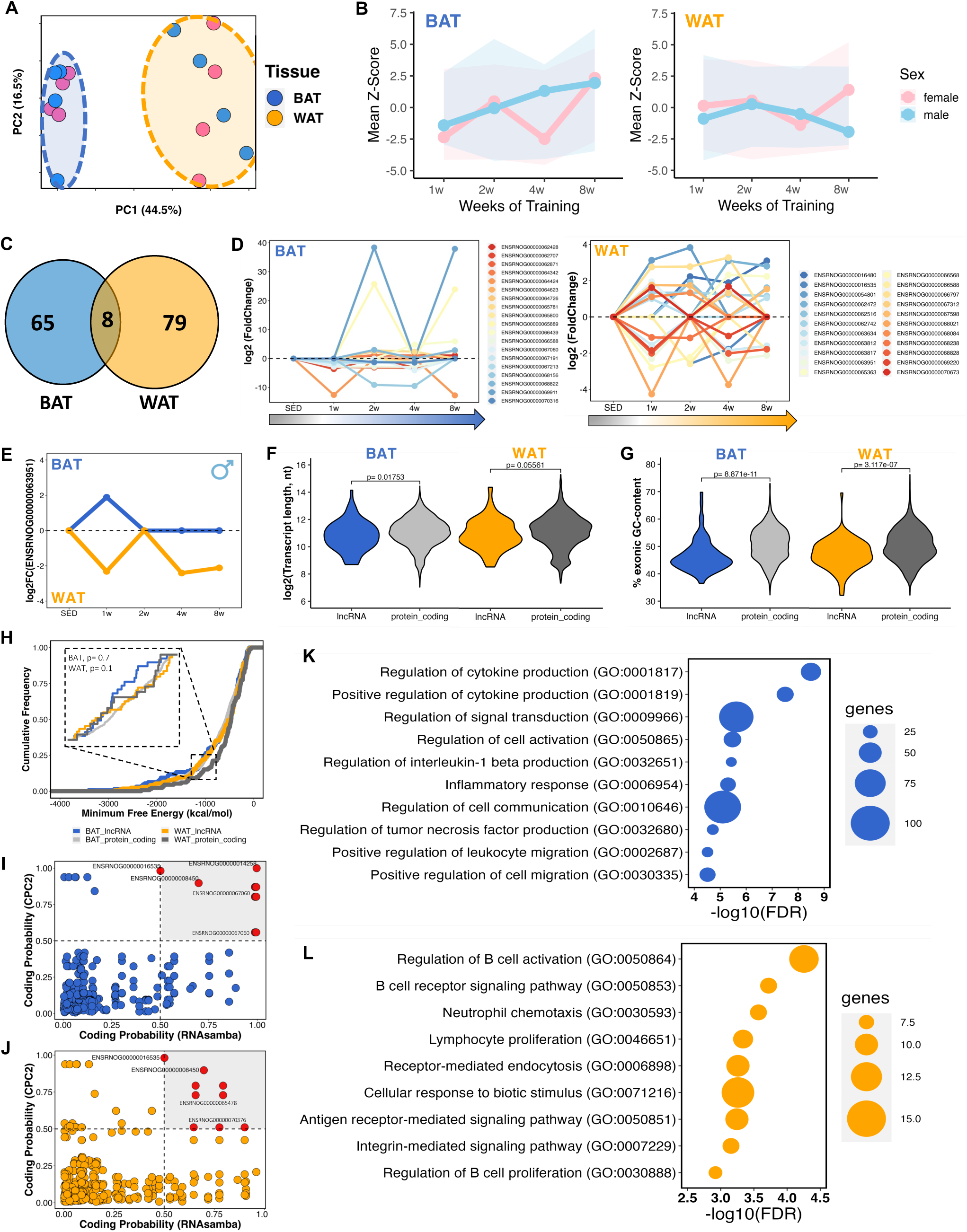
The transcriptional landscape of lncRNAs in adipose tissues after 8 weeks of aerobic exercise. **(A)** Principal Component Analysis (PCA) demonstrating the clustering pattern of all identified lncRNAs in the brown adipose tissue (BAT, blue) and white adipose tissue (WAT, orange) across different training weeks. **(B)** Mean Z-score distribution of all differentially expressed lncRNAs (DELncs) in BAT and WAT during the 8-week training regimen, stratified by sex (male rats in blue, female rats in pink). **(C)** Venn diagram displays common DELncs between BAT and WAT, with the majority of DELncs being unique to each tissue. **(D)** Gene expression changes (log2FC) of DELncs identified across multiple training points in BAT and WAT, respectively. **(E)** Identification of the common DELnc ENSRNOG00000063951 between BAT and WAT, showing dissimilar expression pattern in each tissue across different exercise time points. **(F)** Comparison of the length in nucleotides (nt) of DELncs and differentially expressed protein_coding genes (DEGs) in BAT and WAT. **(G)** Comparison of the exonic GC content percentage in BAT and WAT. **(H)** Minimum Free Energy (MFE) analysis of DELncs and DEGs in BAT and WAT, indicating no statistically significant differences (ns) between lncRNAs and mRNAs. **(I-J)** Scatter plots illustrating protein-coding potential of lncRNAs in BAT (blue) and WAT (orange). **(K-L)** Gene network correlation between DELncs-DEGs revealed specific gene ontology pathways in BAT and WAT, respectively.

One lncRNA (ENSRNOG00000063951) demonstrated opposing expression patterns; upregulated in male BAT and downregulated in male WAT during the initial week of training, with sustained downregulation in WAT at weeks 4 and 8 (**Figure 4E**). This pattern suggests a role for this lncRNA during the early stages of adaptation to training in adipose tissues. Structural analysis confirmed that the differential lncRNAs exhibited significant differences in length (**Figure 4F**) and GC content (**Figure 4G**), while their MFE was comparable to that of mRNAs (DEGs) (**Figure 4H**). Furthermore, lncRNAs exhibited significantly reduced expression compared to protein-coding genes in both adipose tissues (**Figure S9B and S9R**). We also conducted assessments of protein-coding capability by analyzing all transcript sequences, identifying 94% and 95% as non-coding in BAT and WAT, respectively (**Figure 4I-J**).

To investigate potential regulatory associations between differential lncRNAs and mRNAs, we performed correlation analysis in both tissues. Upregulated genes that correlated with differential lncRNAs exhibited similar GO terms for BAT and WAT related to the regulation of immune system processes and response (**Figure S12E and S12F**). Among distinct pathways, we identified regulation of cytokine production, signal transduction, interleukin-1 beta production, inflammatory response, regulation of TNF production, and positive regulation of cell migration in BAT (**Figure 4K**). In WAT, we observed pathways related to regulation of B cell activation, neutrophil chemotaxis, lymphocyte proliferation, receptor-mediated endocytosis, cellular response to biotic stimulus, and integrin-mediated signaling pathway (**Figure 4L**).

### Integrative Profile of lncRNAs and Chromatin Accessibility

LncRNAs are key regulators of chromatin organization and epigenetic gene regulation. They can modulate chromatin structure through their own transcription as well as by recruiting, scaffolding, or displacing chromatin-modifying complexes, thereby promoting either transcriptional activation or repression^51^. To investigate whether any of the training-differential lncRNAs were regulated through chromatin accessibility, we analyzed ATAC-seq data available for eight tissues. For each differential lncRNA, we interrogated a ±500 kb window surrounding its genomic locus for differential ATAC peaks as a proxy for local chromatin accessibility potentially linked to lncRNA expression or its regulatory function.

**Figure 5** illustrates the genome-wide distribution of correlations between lncRNA transcripts and ATAC peaks as a function of distance to the TSS, shown for all detected lncRNA transcripts (left panel) and for differential lncRNAs (right panel). This analysis revealed distinct tissue-specific patterns of chromatin–lncRNA associations, with a strong enrichment near promoter regions. We identified 277 such lncRNA–ATAC peak associations in liver, 124 in WAT, 121 in BAT, 89 in *gastrocnemius*, 74 in lung, 71 in kidney, 13 in heart, and 3 in hippocampus (**Figure 5**), with the top hits highlighted in **Figure S13A-F**.

For each lncRNA that was associated with a differential ATAC peak, we then identified the nearest transcription start site (TSS) for a protein-coding gene and calculated an activity score for the corresponding ATAC peak, defined as the inverse of the genomic distance between the peak summit and that TSS plus one base pair (1 / [distance + 1]) (**Figure 5H**). This metric prioritizes proximal lncRNA–peak–gene associations, thereby enriching for putative cis-regulatory interactions where local chromatin accessibility is most likely to influence transcriptional output.

**Figure 5.**
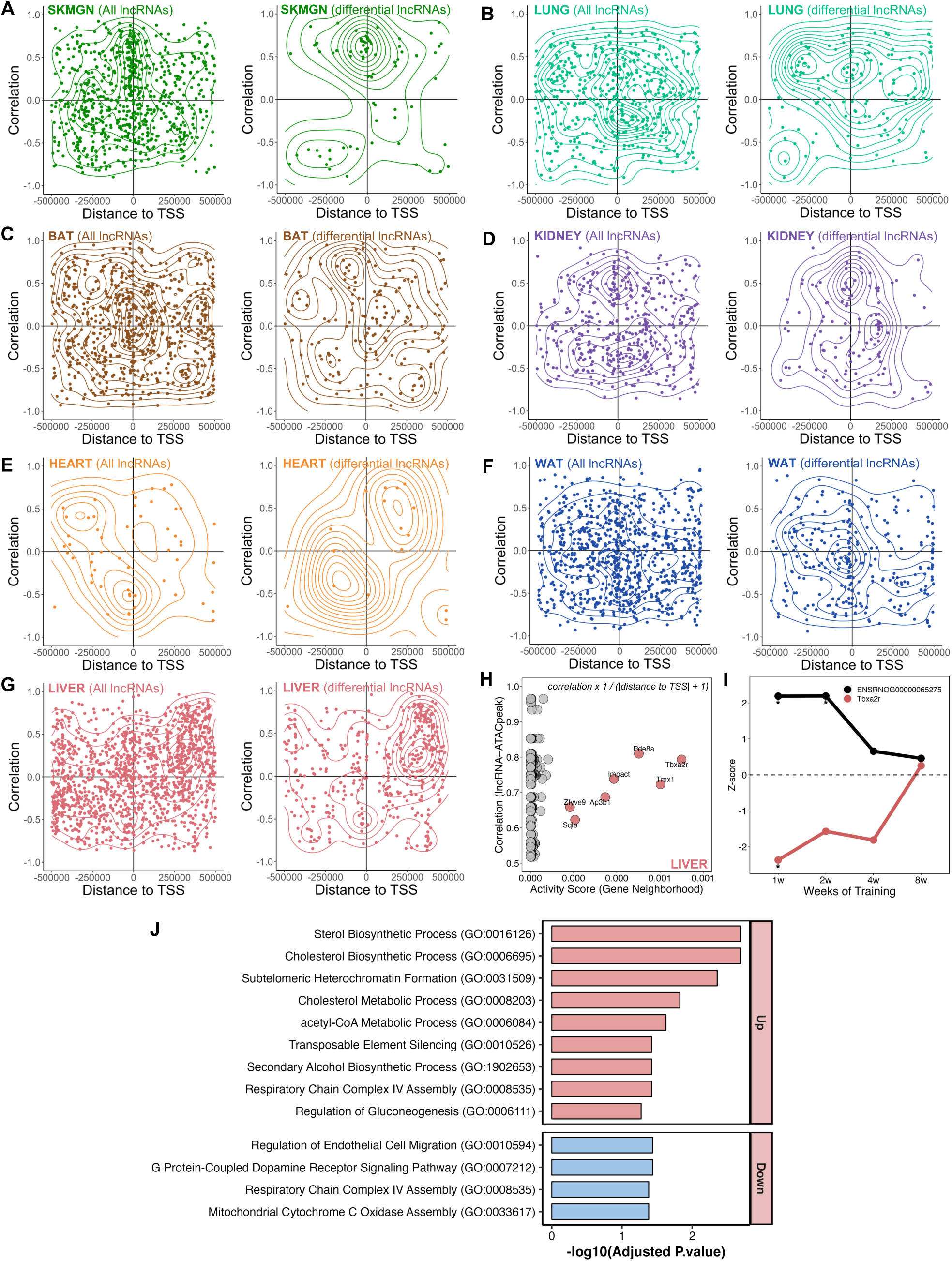
Tissue-specific associations between lncRNAs and chromatin accessibility at regulatory regions. **(A–H)** Scatterplots with density contours showing the correlation between lncRNA expression and ATAC peak accessibility as a function of genomic distance between lncRNA loci and nearby ATAC peaks. Left panels show all expressed lncRNA transcripts; right panels show differentially expressed lncRNAs (FDR < 0.05). Data are shown for **(A)** SKMGN, **(B)** lung, **(C)** BAT, **(D)** kidney, **(E)** heart, **(F)** WAT, and **(G)** liver. **(H)** Dot plot of activity scores in liver, integrating lncRNA–ATAC peak correlation with peak proximity to the TSS of neighboring protein-coding genes. Top-scoring gene neighborhoods include Tbxa2r, Tmx1, and Pde8a. **(I)** Temporal expression trajectories (Z-score) of lncRNA ENSRNOG00000065275 and its associated protein-coding gene Tbxa2r across weeks 1–8 of training in male liver. Asterisks indicate statistically significant differential expression. **(J)** Gene Ontology (GO) enrichment analysis of upregulated and downregulated protein-coding genes identified in lncRNA–ATAC peak–protein-coding triads with positive correlations (*r* ≥ 0.5).

The liver exhibited the largest number of lncRNA–ATAC peak associations. The top-ranked triad was the lncRNA ENSRNOG00000065275 and the Tbxa2r locus. This lncRNA displayed a strong positive correlation with a training-induced ATAC peak located near the Tbxa2r promoter, consistent with enhanced chromatin accessibility (**Figure 5H**). Interestingly, the expression trajectories diverged: while ENSRNOG00000065275 expression increased with training, Tbxa2r mRNA was significantly repressed during the early training weeks (weeks 1 and 2) in males, with partial recovery by week 8 (**Figure 5I**). This inverse relationship between promoter accessibility and gene expression suggests a potential repressive cis-acting mechanism, in which the lncRNA may facilitate chromatin remodeling that uncouples accessibility from productive transcription.

To further explore the potential functional consequences of lncRNA–chromatin interactions, we analyzed lncRNA–ATAC peak–protein-coding gene triads that exhibited a positive correlation between lncRNA expression and peak accessibility in liver (**Figure 5H**). For each triad, we classified the associated protein-coding gene as either upregulated or downregulated in response to exercise. Upregulated genes were significantly enriched in pathways related to sterol and cholesterol biosynthesis, acetyl-CoA metabolism, and chromatin regulation (**Figure 5J**)—all processes closely linked to metabolic remodeling during exercise. In contrast, downregulated genes were enriched for pathways associated with endothelial cell migration, G protein–coupled receptor signaling, and mitochondrial respiratory complex assembly, reflecting vascular and mitochondrial adaptations (**Figure 5J**). These results support the hypothesis that exercise-responsive lncRNAs associated with local chromatin accessibility may contribute to the regulation of distinct transcriptional programs, potentially exerting cis-regulatory control over neighboring protein-coding genes. Additional triads exhibiting either concordant or discordant accessibility–expression dynamics were identified across other tissues (**Figure S13G–L**), underscoring the complexity of lncRNA-mediated chromatin regulation in exercise adaptation.

## DISCUSSION

We leveraged and enhanced the comprehensive multi-omics datasets (transcriptomics, epigenomics, and proteomics) generated by MoTrPAC^23^ to gain a deeper understanding of the lncRNA landscape associated with adaptations induced by progressive endurance training. Our work shows remodeling of the lncRNA landscape, particularly affecting the adrenal glands, BAT, WAT, and the hypothalamus. The changes were remarkably tissue-and sex-specific, with several lncRNAs regulated by chromatin accessibility. Sex differential expression of lncRNAs align with recent findings by Hanks et al.^29^ demonstrating that lncRNAs are significantly more likely to exhibit sex-biased expression than protein-coding genes in human skeletal muscle fiber nuclei, suggesting that noncoding RNAs play a central role in the regulation of sex-specific gene expression, even in the absence of external stimuli^29^. Our data extend these observations by demonstrating that lncRNA expression is not only inherently sex-biased but also dynamically modulated in response to physiological challenges such as exercise.

Tissues directly involved in exercise, such as the skeletal muscles and the heart, were expected to exhibit a higher abundance of differentially expressed lncRNAs. Interestingly, we found that the tissues with greatest regulation of lncRNAs were the hypothalamus, adipose tissues (BAT and WAT), and adrenal glands (**Figure 1**). The hypothalamus is the primary tissue responsible for the control of whole-body homeostasis. Through hormonal and metabolic regulation, it modulates food intake, thirst, temperature, blood pressure, heart rate, and the secretion of various essential hormones^52–54^. We observed multiple differential lncRNAs in the hypothalamus of trained animals, with the first four weeks showing greater and more consistent regulation (**Figure 1**). Thus, these lncRNAs may have fundamental roles in adaptation to exercise and homeostasis functions regulated by the hypothalamus. However, our understanding of mRNA and lncRNA expression and function in the hypothalamus–particularly in response to physiological stimuli-remains limited. For example, Katz et al.^55^ demonstrated that both the hypothalamus and skeletal muscle (*gastrocnemius*) display similar metabolic responses to exercise. Both tissues activate the AMPK pathway following voluntary wheel running in mice, stimulating ATP production and mitochondrial biogenesis, yet they diverge in the activation pattern of the AKT/mTOR pathways (promotes cell growth and protein synthesis), which depend on the mode and intensity of exercise^55^. Regarding lncRNAs, other studies have been examining hypothalamic lncRNA expression in response to other physiological stimuli, such as food deprivation or heat stress in mice^56^, rats^57^, chickens^58^, sheep^59^, goats^60^, pigs^61^, and cows^62^.

Adipose tissues (BAT and WAT) are critical for the regulation of whole-body metabolism, as well as for maintaining homeostasis in response to various stimuli^63^. There are abundant molecular changes in adipose tissues in diseases such as obesity and type 2 diabetes, as well as in response to exercise. These changes involve the regulation of fatty acids, energy and lipid metabolism, hormonal and enzymatic activity, expression and secretion of adipokines (e.g., Apelin and Adiponectin), and immune, angiogenic, and fibrotic remodeling^23,47,64,65^. Furthermore, the beneficial effects of physical exercise on the modulation of adipose tissue are well known, highlighting positive changes in immune and inflammatory responses, as well as lipid metabolism^23,66–69^. Our work aligns with these and other studies, as we document the possible regulation of pathways related to the immune response in both adipose tissues by lncRNAs (**Figure 4**).

Several lncRNAs have important functions in adipogenesis and in the maintenance of adipose tissue^70–73^. However, only a few have been reported to be activated or suppressed after physical exercise and to have direct roles in adipose tissue physiology^74,75^. For example, Wu et al.^74^ showed that 8 weeks of endurance exercise inhibited the expression of the lncRNA SRA in obese mice. This inhibition could trigger the activation of the p38/JNK pathway, suppressing the expression of PPARψ and its downstream target genes, resulting in improved lipid metabolism.

Although adipocytes are the primary cells in adipose tissue, it also contains immune cells, endothelial cells, fibroblasts, stem cells and progenitors, all essential for tissue maintenance and function^76^. For instance, Yang et al.^77^ conducted an extensive single-cell mapping of adipose tissue and skeletal muscle, underscoring the significance of mesenchymal stem cells (MSCs) in tissue adaptations linked to obesity and exercise in mice. They found that MSCs play a role in regulating pathways related to extracellular matrix remodeling and circadian rhythm in response to high-fat diet and exercise^77^. There is compelling evidence showing that adipose MSCs undergo substantial transcriptional and post-transcriptional regulation during adipogenic differentiation^78^. This regulation encompasses alterations in the expression of both mRNAs and lncRNAs^78,79^. Furthermore, multiple ongoing studies are exploring the regulatory functions of lncRNAs in obesity progression, adipogenesis, and the influence of physical activity. For example, Martinez et al.^80^ reported the presence of numerous uncharacterized small open reading frames (smORFs) within both protein-coding genes and lncRNAs in mouse adipocytes. These smORFs encode microproteins, including one that exhibits orexigenic activity in obese mice^80^. In our study, we were unable to identify bona-fide microproteins from the mass-spectrometry data. Importantly, the identification of fast-turnover and low-abundance proteins, including microproteins, by mass spectrometry remains inherently challenging due to a range of technical complexities. These include sample composition, low expression levels, microprotein translation dynamics (i.e., ribosome and polysome association), protein half-life, among other factors^81–84^.

The current work emphasizes the importance of analyzing different molecular layers affected by exercise. It highlights the differential expression and regulatory role of lncRNAs at various levels — DNA (genomic/epigenomic), RNA (transcriptional/post-transcriptional), and protein (translation/protein-binding). We anticipate that these findings will enable hypothesis-driven exploration of lncRNAs in response to exercise and training. The identification of lncRNA regulation across multiple tissues in response to endurance training demonstrates the complexity of exercise-induced molecular adaptations and establishes a foundation for understanding the regulation by noncoding RNA in a physiological context. The marked sex-specific differences further underscore the need to incorporate both sexes in future multi-omics studies to fully capture the functional roles of lncRNAs that likely contribute to the many health benefits of exercise. Collectively, by establishing the first comprehensive atlas of exercise-responsive lncRNAs across multiple tissues and timepoints, we provide a critical resource for the exercise genomics community and highlight the untapped regulatory potential of the non-coding transcriptome in human health and disease.

### Limitations of the study

Despite the comprehensive exploration of lncRNA responses to endurance exercise training in male and female rats, the study has limitations. The exclusive focus on rats requires caution when extrapolating the findings to humans, given species-specific variations, lack of gene conservation, and variations in physiological responses. The study emphasizes long-term responses to endurance training, with samples collected 48 hours after the last exercise bout and therefore does not capture the full dynamics of the short-term responses. In addition, the different time points are represented by unique rats, increasing the potential variability for lowly expressed transcripts such as lncRNAs. The interpretation of tissue-and sex-specific lncRNA expression patterns remains challenging, since differences in cellular composition, hormonal environment, and inter-organ communication can profoundly influence the observed responses. Finally, the functional significance of identified lncRNAs is incompletely understood, underscoring the need for future studies to clarify their roles in exercise adaptation.

## MATERIALS AND METHODS

### Animal and Endurance Training Protocol

The animal procedures, endurance training interventions, and primary molecular data generation used in this study have been described previously^23^. In brief, male and female Fischer 344 rats were procured from the National Institute of Aging (NIA) rodent colony. These rats were accommodated in pairs under a reverse dark-light cycle, with environmental conditions maintained at a temperature of 20-25° C. They were provided with standard chow (Lab Diet 5L79) as their diet. After an initial period of acclimatization, the rats were randomly assigned to either the training or control groups. The rats were categorized into three cohorts: 8-week rats, which were further divided into training and control groups; 4-week rats, all designated for training; and 1-and 2-week rats, which were randomly assigned to either 1 or 2 weeks of training. A total of 50 rats (comprising 5 males and 5 females per time point) were included in molecular analyses for all -omics except proteomics which was assayed on 60 rats (6 males and 6 females per timepoint). All rats started training at 6 months of age, utilizing a Panlab 5-lane rat treadmill (Harvard Instruments, Model LE8710RTS). The training regimen consisted of 5 days of exercise per week, employing a progressive protocol designed to maintain an intensity equivalent to approximately 70% of VO_2_max, which involved incremental adjustments in grade and speed. During the last two weeks of training, the maximum duration of exercise sessions was set at 50 minutes. The initial treadmill speed was determined based on VO_2_max measurements obtained after the familiarization period. All animal procedures were conducted during the dark cycle and received approval from the Institutional Animal Care and Use Committee at the University of Iowa.

### Collection and Processing of Tissues

All tissue samples were harvested 48 hours following the final exercise session. A fasting period of three hours preceded the dissection process. During dissection, rats were anesthetized with inhaled isoflurane (1-2 %) and remained under anesthesia until expiration. Cardiac puncture was used to obtain blood, after which the *gastrocnemius* muscle, subcutaneous white adipose, right lobe of the liver, heart, and lungs were sequentially extracted. Removal of the heart led to fatality. Subsequently, decapitation was performed using a guillotine, followed by the removal of the brain, including the hypothalamus, right and left hippocampus, right and left cerebral cortex. Following decapitation, the right kidney, right and left adrenal glands, spleen, brown adipose tissue, small intestine, colon, right testes or ovaries, and right *vastus lateralis* were excised, in this particular order. All collected tissue specimens were promptly flash-frozen in liquid nitrogen and stored at -80° C.

### Multi-omics Data Generation and Processing: Transcriptomics

Full details of the methods used for sample processing, data collection, processing, normalization, and batch correction for transcriptomics (RNA sequencing, RNA-seq) and epigenomics (Assay for Transposase-Accessible Chromatin using sequencing, ATAC-seq) platforms are described elsewhere^23^. In brief, RNA-seq data were processed using the previously described pipeline^23^, with the main difference being alignment to the Ensembl Rattus norvegicus mRatBN7.2 genome (release 108). The assembly was performed using STAR (v.2.7.0d) and gene quantification with RSEM (v.1.3.1). Filtering and normalization were performed separately by tissue, retaining genes with >0.5 counts per million in at least two samples. The openWDL-based implementation of the RNA-seq pipeline on Google Cloud Platform is available at https://github.com/MoTrPAC/motrpac-rna-seq-pipeline.

Differential expression analysis was performed independently for each tissue using DESeq2 on raw counts, applying its median-of-ratios normalization method. For exploratory analysis, filtered raw counts were normalized using the trimmed mean of M-values method (TMM; edgeR::calcNormFactors) and transformed to log counts per million (logCPM; edgeR::cpm). Male and female samples were analyzed separately, with contrasts comparing each training timepoint to the corresponding sex-matched sedentary controls to estimate time- and sex-specific effect sizes. Covariates included RNA integrity number (RIN), median 5′–3′ bias, percent of reads mapping to globin genes, and percent PCR duplicates (quantified using Unique Molecular Identifiers, UMIs).

To investigate lncRNA expression patterns between the untrained (i.e., sedentary) and trained (i.e., endurance exercise) cohorts, we first filtered the gene set to include only transcripts with a mean Fragments Per Kilobase of transcript per Million mapped reads (FPKM) ≥ 1, thereby focusing on adequately expressed genes. Differentially expressed lncRNAs (DELncs) were then identified using a *p-value* threshold of ≤ 0.01 and an absolute log2 fold change (|log2FC|) ≥ 1, a strategy chosen to enhance sensitivity toward low-abundance and variable lncRNAs and to avoid excluding relevant candidates. In parallel, we also performed a stringent exploratory analysis using multiple-testing correction (10% Benjamin-Hochberg FDR adjustment, log2 fold change (|log2FC|) ≥ 0.5), which yielded a total of 433 unique differential lncRNAs across the 18 tissues analyzed. Together, these complementary approaches allowed us to balance sensitivity in detecting dynamic lncRNA signals with specificity in establishing a high-confidence set of candidates for downstream analyses, a strategy commonly adopted in lncRNA studies given their typically low abundance and high variability^85–88^.

### Multi-omics Data Generation and Processing: Epigenomics

For ATAC-seq, data processing largely followed our previously published workflow^23^, with modifications specific to this study. Briefly, nuclei were isolated from tissue samples using a modified Omni-ATAC protocol. Following DAPI staining and counting, 50,000 nuclei were subjected to transposition, purified, and PCR-amplified. Libraries were sequenced (50bp paired-end) on an Illumina NovaSeq 6000, targeting ∼35 million read pairs per sample. Sequencing reads were demultiplexed and processed using the ENCODE ATAC-seq pipeline (v.1.7.0) (https://github.com/ENCODE-DCC/atac-seq-pipeline). Notably, in this study reads were aligned to the updated Rattus norvegicus mRatBN7.2 (Rn7) genome using Bowtie2, and all downstream analyses described here are novel. Peak calling was performed using MACS2 (v.2.2.4) in replicate mode combining all biological replicates per time point and sex. Optimal peaks from all tissues were concatenated, trimmed to 200 base pairs around the summit, and merged to generate a master peak list. This master peak list was intersected with the filtered alignments from each sample using bedtools coverage (options -nonamecheck and -counts) to generate a peak by sample matrix of raw counts. The remaining steps such as outlier removal, normalization and differential analysis were applied separately on raw counts from each tissue. Only peaks in autosomes and sex chromosomes and those with at least 10 reads in 4 or more samples were included in downstream analysis.

Filtered raw counts were normalized with DESeq2 for each tissue separately. Batch effect was modeled by including the *fraction of reads in peaks*, and sample processing batch (coded as a factor), in the design formula. Contrasts compared each training timepoint to the corresponding sex-matched sedentary controls to estimate time- and sex-specific effect sizes. To identify peaks that were significantly altered at any time point, we used an F-test implemented in limma. For each feature, male- and female-specific *p-values* were combined using Fisher’s sum-of-logs meta-analysis, yielding a single *p-value*, referred to as the training p-value. These values were adjusted across all datasets within each ‘ome’ using Independent Hypothesis Weighting (IHW) with tissue as a covariate. Training-differential features were defined at an IHW-adjusted FDR ≤ 0.05. This stricter multiple-testing correction was chosen for ATAC-seq to ensure high-confidence peak detection, given its role as the structural reference for downstream lncRNA–chromatin accessibility integration analyses.

### ATAC-seq analysis of lncRNA-associated peaks

To analyze shared patterns of exercise training response in the lncRNA and ATAC-seq data, we first needed to annotate each lncRNA and ATAC peak with its genomic location. We used the BioMart function *getBM()* to annotate the lncRNAs and the ChipSeeker function *annotatePeak()* to annotate the ATAC-seq peaks. We measured the distance between the midpoint of each lncRNA locus and ATAC-seq peak in the genome and identified local lncRNA-peak pairs that are located within 500kb of each other on a given chromosome.

We restricted the analysis to local pairs in which the ATAC-seq peak showed a significant training response (IHW-adjusted FDR < 0.05 from the F-test). For each such pair, log2FC values from all training timepoints were used to compute the Pearson correlation coefficient (𝑟) between the lncRNA and ATAC-seq peak responses. For each tissue, we plotted the relationship between genomic distance and exercise response correlation for each local pair and estimated the 2D kernel density of this relationship by the function *geom_density_2d()* for each tissue and at different thresholds of *adjusted p-value* cutoff for the significance of the lncRNA exercise training response.

To prioritize candidate regulatory lncRNAs, we computed an activity score for each lncRNA–ATAC peak pair and its nearest protein-coding gene as:

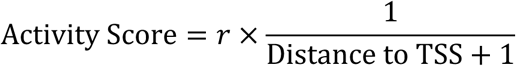

where *r* is the Pearson correlation coefficient, and “Distance to TSS” is the absolute distance in base pairs between the ATAC-seq peak midpoint and the transcription start site (TSS) of the nearest protein-coding gene. This metric favors high-correlation associations proximal to regulatory regions, suggesting potential roles in local cis regulation of chromatin.

To explore the functional consequences of lncRNA–chromatin interactions, we focused on lncRNA–ATAC peak–protein-coding gene triads that showed a positive correlation between lncRNA expression and peak accessibility (*r* ≥ 0.5). When multiple triads mapped to the same gene, only the one with the highest activity score was retained. Based on RNA-seq data, the associated protein-coding genes were classified as upregulated or downregulated in response to exercise. Up- and downregulated gene sets were then analyzed separately for Gene Ontology enrichment using EnrichR^89^; significantly enriched terms (FDR < 0.05) were reported.

### Multi-omics Data Generation and Processing: LC-MS/MS Proteomics

The methods used to generate and process the proteomics data are described in detail elsewhere^23^. Briefly, seven distinct rat tissues were subjected to lysis, protein extraction, and enzymatic digestion. The resulting peptides were labeled with TMT11-plex reagents for relative quantification. Samples were then fractionated using offline basic reversed-phase chromatography and analyzed by liquid chromatography-tandem mass spectrometry (LC-MS/MS). The raw mass spectrometry data were processed using the exact same custom cloud-based MS-GF+ pipeline as in our previous publication^23^, with two key updates: (**i**) the pipeline was uniformly applied across all seven tissues, and (**ii**) database searches were performed against an updated rat reference proteome based on the Rn7 genome. This reference was expanded to include 159 lncRNAs that were either computationally predicted or annotated (i.e., RGD) as protein-coding, with all putative open reading frames (ORFs) translated in the three frames and incorporated into the search database to enable unbiased detection of non-canonical microproteins. Subsequent data processing, including TMT reporter ion quantification, normalization, and differential abundance analysis, also followed the previously established methods.

### Prediction of lncRNA Features

The minimum free energy (MFE) associated with secondary structure formation in lncRNAs was predicted using RNAfold software (v.2.4.14)^90^. This prediction relied on the nearest-neighbor thermodynamic model and utilized a FASTA input file containing selected lncRNA and protein-coding sequences from Ensembl (mRN7.2). To statistically compare lncRNA and protein-coding groups, transcript length and GC content were evaluated using the Mann–Whitney U test, whereas MFE distributions were compared using the Kolmogorov–Smirnov test. All tests were performed within each tissue separately, with *p* < 0.05 considered statistically significant.

Coding potential probability was assessed using CPC2^42^ and RNAsamba^43^ software. We defined cutoff values for coding potential as > 0.5 for CPC2 and > 0.5 for RNAsamba. Additionally, we conducted Basic Local Alignment Search Tool (BLAST, available at https://blast.ncbi.nlm.nih.gov/Blast.cgi) search to identify orthologous sequences across species or microprotein sequence, providing insights into conserved regions within the lncRNAs.

To identify small open reading frames (smORFs) within lncRNAs, we utilized the Open Reading Frame Finder (NCBI ORFfinder, available at https://www.ncbi.nlm.nih.gov/orffinder/). The search was restricted to ORFs with a minimum length of 30 nucleotides and an ATG start codon (-ml 30 -s ATG). Input sequences were provided in FASTA format.

Interactions between lncRNAs and microRNAs were predicted using AnnoLnc2^91^, which computes conservation scores within primate, mammal, and vertebrate clades to increase prediction confidence. For each identified miRNA, predicted target mRNAs were retrieved from miRDB^92^. A lncRNA–miRNA–mRNA interaction network was generated by cross-referencing the predicted target mRNAs with the set of upregulated differentially expressed protein-coding genes in the target tissues. Gene Ontology (GO) analysis was performed on these genes using STRING database^93^. The resulting networks were visualized using Cytoscape (v.3.5.0).

### Data and code availability

MoTrPAC data are publicly available via http://motrpac-data.org/data-access. For data-related inquiries, please contact the MoTrPAC Help Desk at. Additional resources can be found at http://motrpac.org and https://motrpac-data.org/. RNA-seq and ATAC-seq raw data were deposited at the Gene Expression Omnibus (GEO) under accession GSE242358; and proteomics data were deposited at MassIVE under accessions MSV000092911 and MSV000092922. MoTrPAC data processing pipelines for RNA-seq, ATAC-seq, RRBS and proteomics are available in the following GitHub repositories: https://github.com/MoTrPAC/motrpac-rna-seq-pipeline, https://github.com/MoTrPAC/motrpac-atac-seq-pipeline, https://github.com/MoTrPAC/motrpac-rrbs-pipeline, and https://github.com/MoTrPAC/motrpac-proteomics-pipeline.

## Funding

This study was supported by MoTrPAC NIH grants U24OD026629 (Bioinformatics Center), U24DK112349, U24DK112340, U24DK112331, U24DK112348 (Chemical Analysis Sites), U24AR071113 (Consortium Coordinating Center) and U01AG055133 (PASS/Animal Site). The work was further supported by the Wu Tsai Human Performance Alliance at Stanford University and the Joe and Clara Tsai Foundation.

**Supplementary Figure S1.** Expression and dynamics of differentially expressed lncRNAs (DELncs) across various tissues during an 8-week training regimen.

**Supplementary Figure S2.** Sex-specific differentially expressed lncRNAs (DELncs) in various tissues during an 8-week training regimen.

**Supplementary Figure S3.** Multidimensional analysis of tissue samples from male and female rats following exercise training.

**Supplementary Figure S4.** Chromosomal distribution of lncRNAs identified after aerobic exercise training across 18 rat tissues.

**Supplementary Figure S5.** Analysis of orthologs between rat lncRNAs and mouse and human transcripts.

**Supplementary Figure S6.** Expression of predicted lncRNAs annotated as protein-coding genes in RGD.

**Supplementary Figure S7.** Comparative analysis of the rat lncRNA ENSRNOG00000064728 and its ortholog ENSMUSG00000085837 in mice.

**Supplementary Figure S8.** Muscle-specific variation in lncRNA expression during an 8-week training regimen.

**Supplementary Figure S9.** Comparative expression analyses of lncRNAs and protein-coding genes across 18 rat tissues following exercise.

**Supplementary Figure s10.** Comparative analysis of lncRNA and protein-coding transcripts across various tissues.

**Supplementary Figure s11.** Comparative analysis of coding probability among differentially expressed lncRNA isoforms across tissues.

**Supplementary Figure s12.** Adipose tissue-specific variation in lncRNA expression during an 8-week training regimen.

**Supplementary Figure s13.** Tissue-specific lncRNA–chromatin accessibility correlations and candidate cis-regulatory interactions.

