## Supplementary figures for "The Long Non-coding RNA Landscape of Endurance Exercise Training"

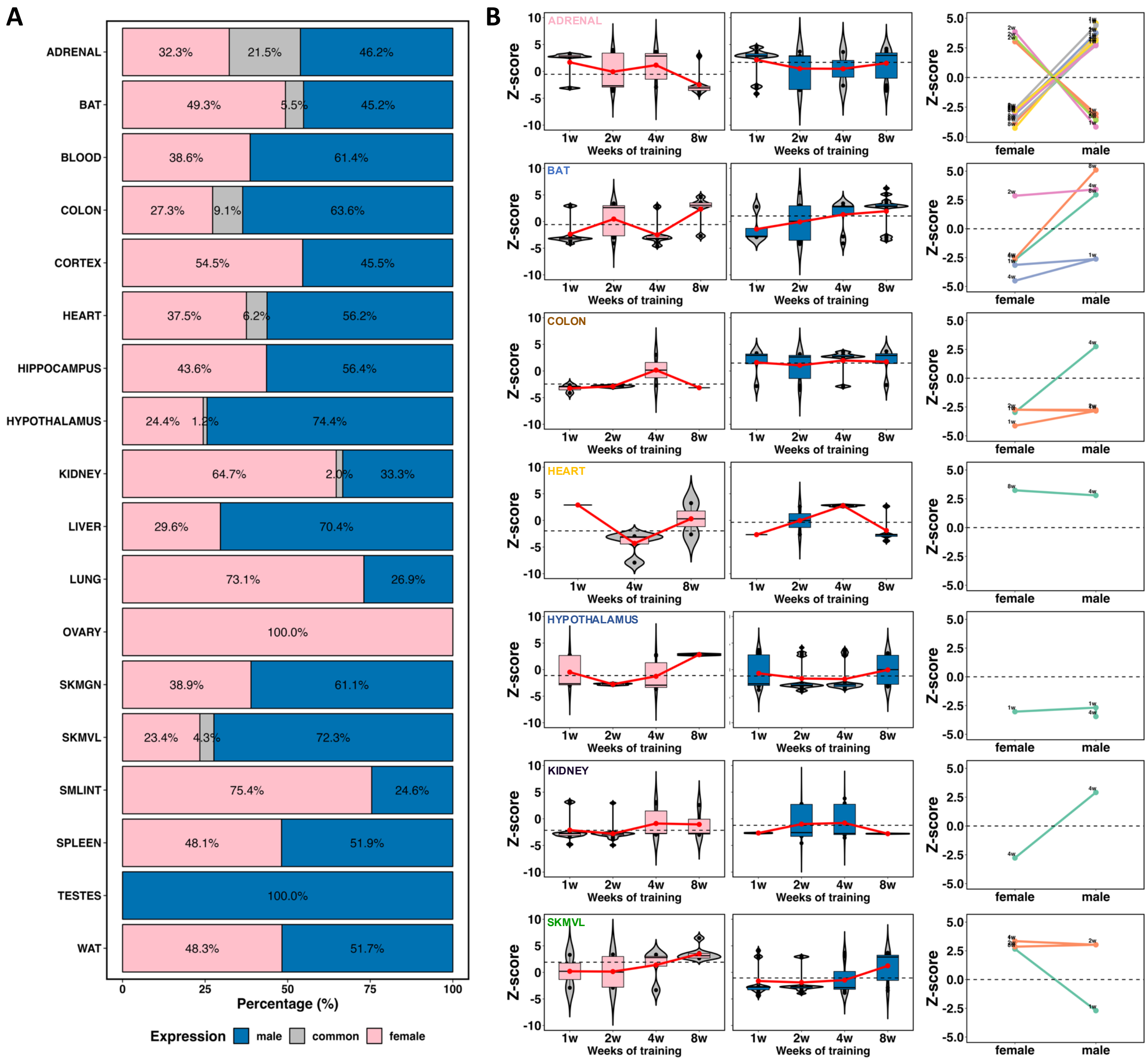


**Figure S1**. **Expression and dynamics of differentially expressed lncRNAs (DELncs) across various tissues during an 8-week training regimen. (A)** Proportion of DELncs expressed in females (pink) and males (blue), and those common to both sexes (gray) across tissues. Only seven tissues exhibited DELncs common to both sexes. **(B)** Temporal expression patterns of common DELncs in representative tissues (adrenal, BAT, colon, heart, hypothalamus, kidney, and SKMVL). Boxplots and violin plots display Z-scores for common DELncs expression, while line plots (right panel) summarize overall Z-score trends by sex, highlighting distinct expression dynamics in specific tissues and time points.

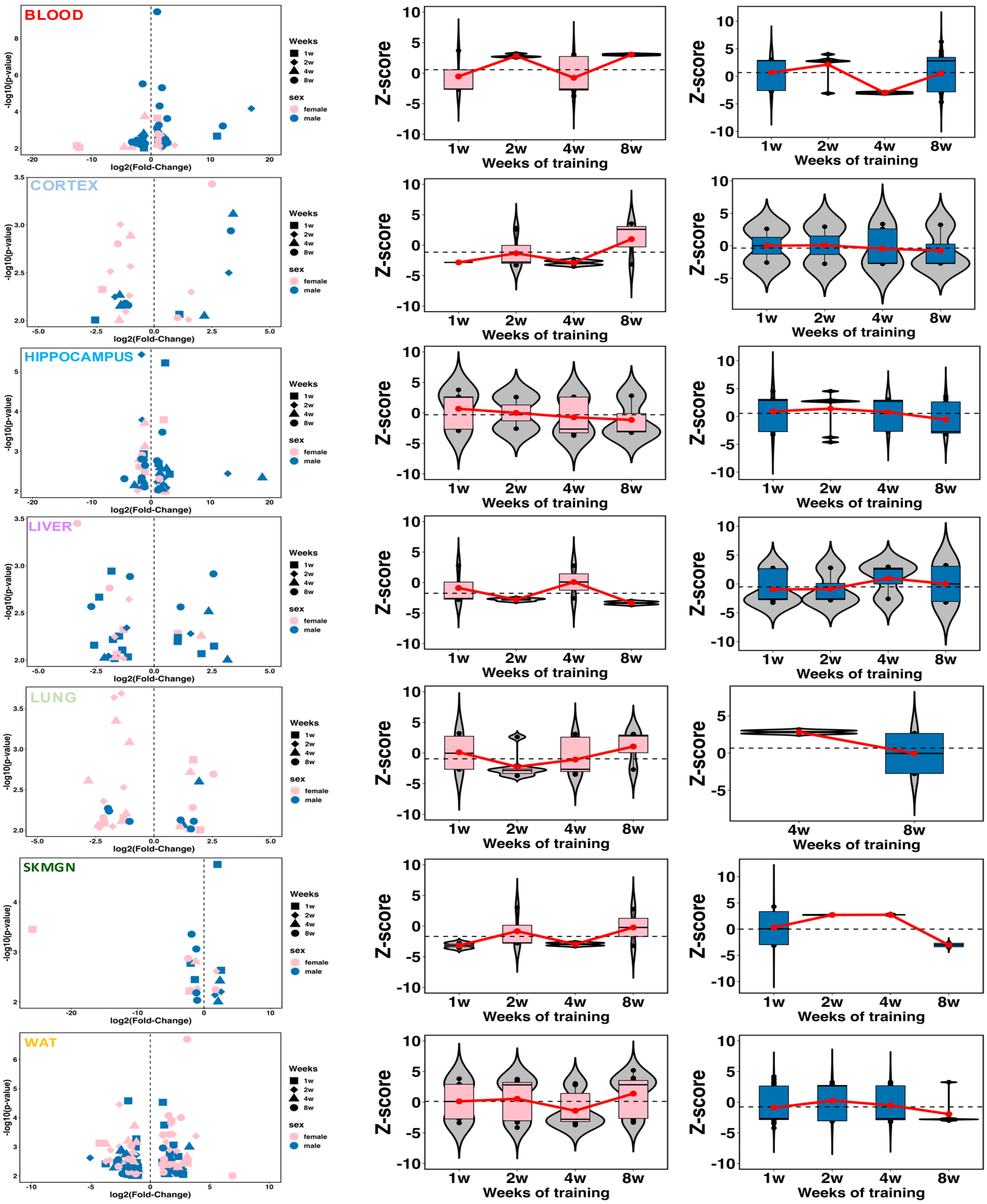


**Figure S2**. **Sex-specific differentially expressed lncRNAs (DELncs) in various tissues during an 8-week training regimen.** Volcano plots showing sex-specific lncRNAs expressed in blood, cortex, hippocampus, liver, lung, SKMGN, and WAT. Pink and blue points represent differentially expressed lncRNAs (DELncs) specific to females and males, respectively (left panel). Boxplots and violin plots illustrating Z-score distributions of DELncs expression stratified by sex (pink for females, blue for males).

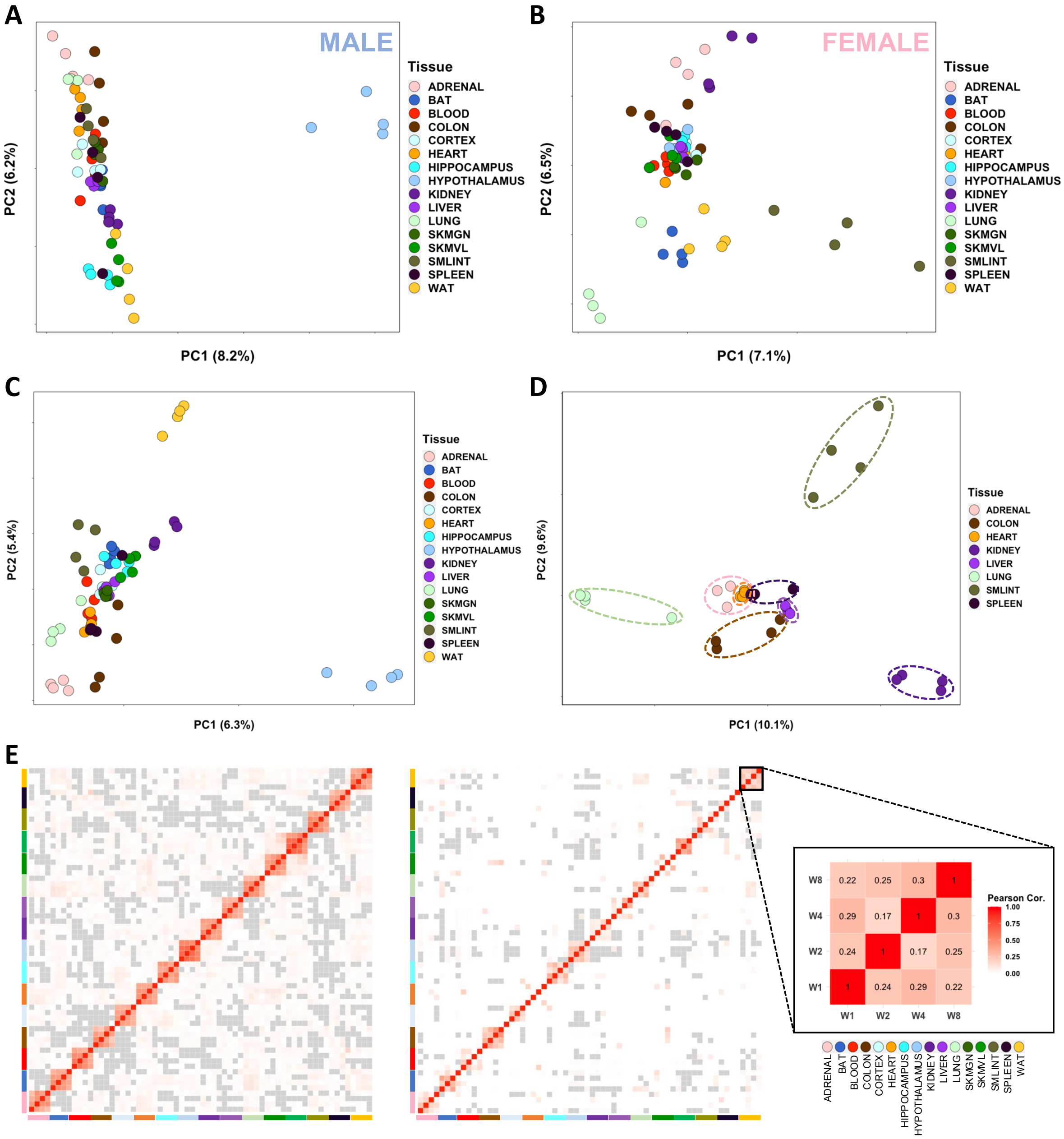


**Figure S3**. **Multidimensional analysis of tissue samples from male and female rats following exercise training. (A-B)** Principal Component Analysis (PCA) of tissue samples from male (**A**) and female (**B**) rats, highlighting distinct clustering patterns across tissues. **(C)** PCA combining all lncRNAs (male and female) and tissue samples collected after 1, 2, 4, and 8 weeks of exercise training, showing overall transcriptional variation. **(D)** PCA analysis focusing on specific internal tissues (e.g., adrenal, heart, kidney, liver, lung, small intestine, spleen) to highlight tissue-specific clustering. **(E)** Heatmap of Pearson correlation coefficients comparing expression patterns of all lncRNAs (left panel) and differentially expressed lncRNAs (DELncs) (right panel) across training weeks. The inset in (**E**) provides detailed Pearson correlation values between weeks for specific tissues.

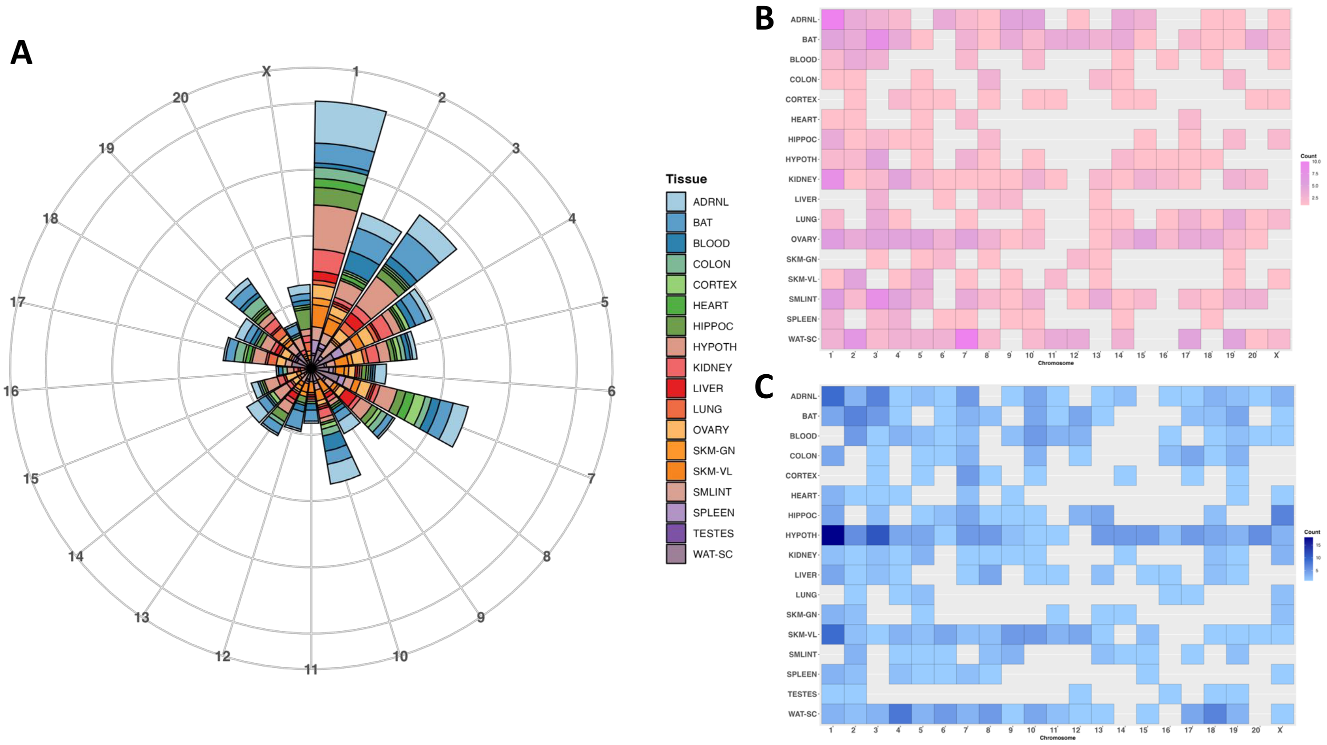


**Figure S4**. **Chromosomal distribution of lncRNAs identified after endurance exercise training across 18 rat tissues. (A)** Circular bar plot illustrating the chromosomal distribution of lncRNAs across all analyzed tissues. Each bar represents a specific chromosome, with colors indicating tissue types. **(B-C)** Heatmaps showing the chromosomal distribution of lncRNAs stratified by sex: (**B**) female rats (pink) and (**C**) male rats (blue). The intensity of the color represents the number of lncRNAs identified in each tissue-chromosome combination.

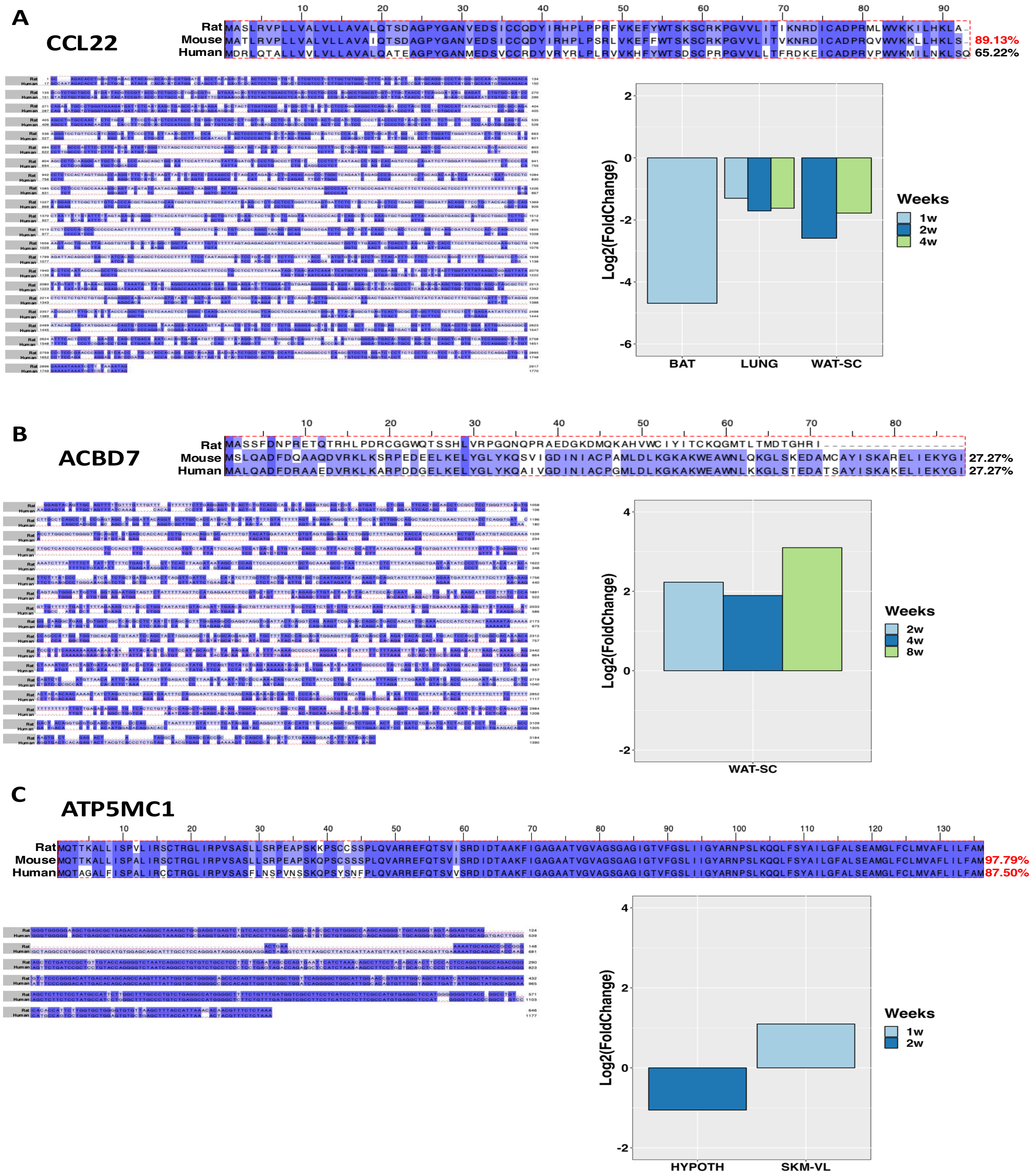


**Figure S5. Analysis of orthologs between rat lncRNAs and mouse and human transcripts. (A-C)** The top panels show the predicted ORF of the protein-coding sequence and its conservation percentage among rats, mice, and humans. The bottom left panels depict RNA sequence conservation between rat and human transcripts for Ccl22 (**A**), Acbd7 (**B**), and Atp5mc1 (**C**). The bottom right panels display expression levels in the respective tissues after 8 weeks of physical activity.

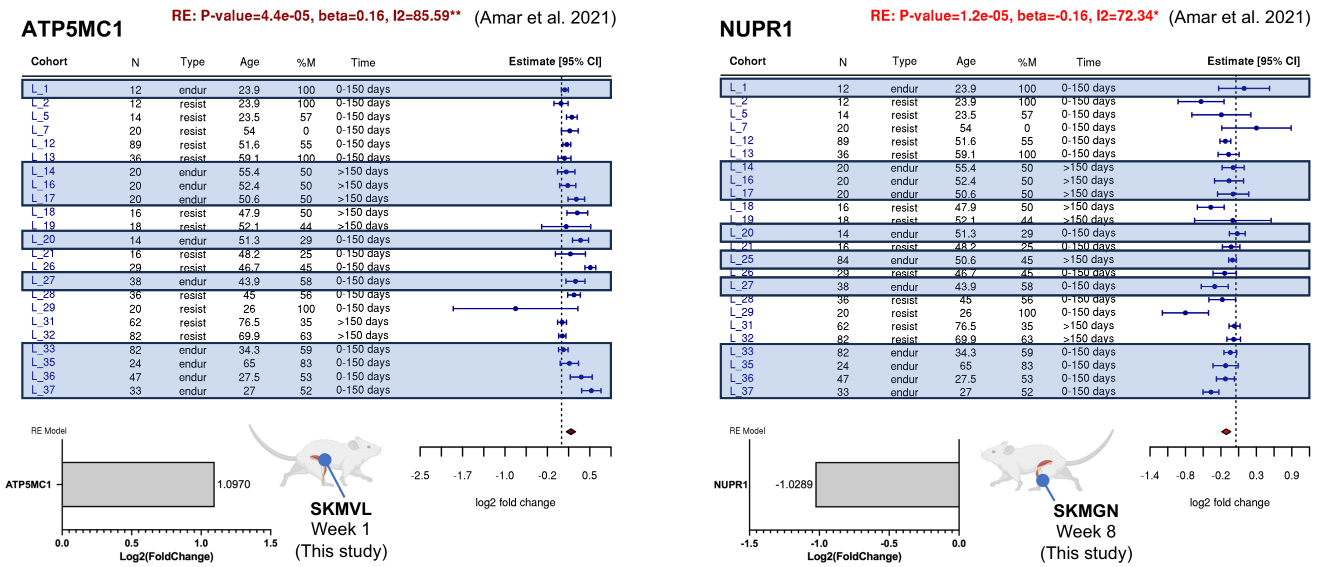


**Figure S6. Expression of predicted lncRNAs annotated as protein-coding genes in RGD.** Shown are two transcripts (Atp5mc1 and Nupr1) predicted as lncRNAs in Ensembl but annotated as protein-coding genes in the Rat Genome Database (RGD). Forest plots display longitudinal human skeletal muscle gene expression data from Amar et al. (2021), with corresponding expression changes from this study shown for SKMVL (Week 1) and SKMGN (Week 8). Blue rectangles highlight endurance exercise cohorts.

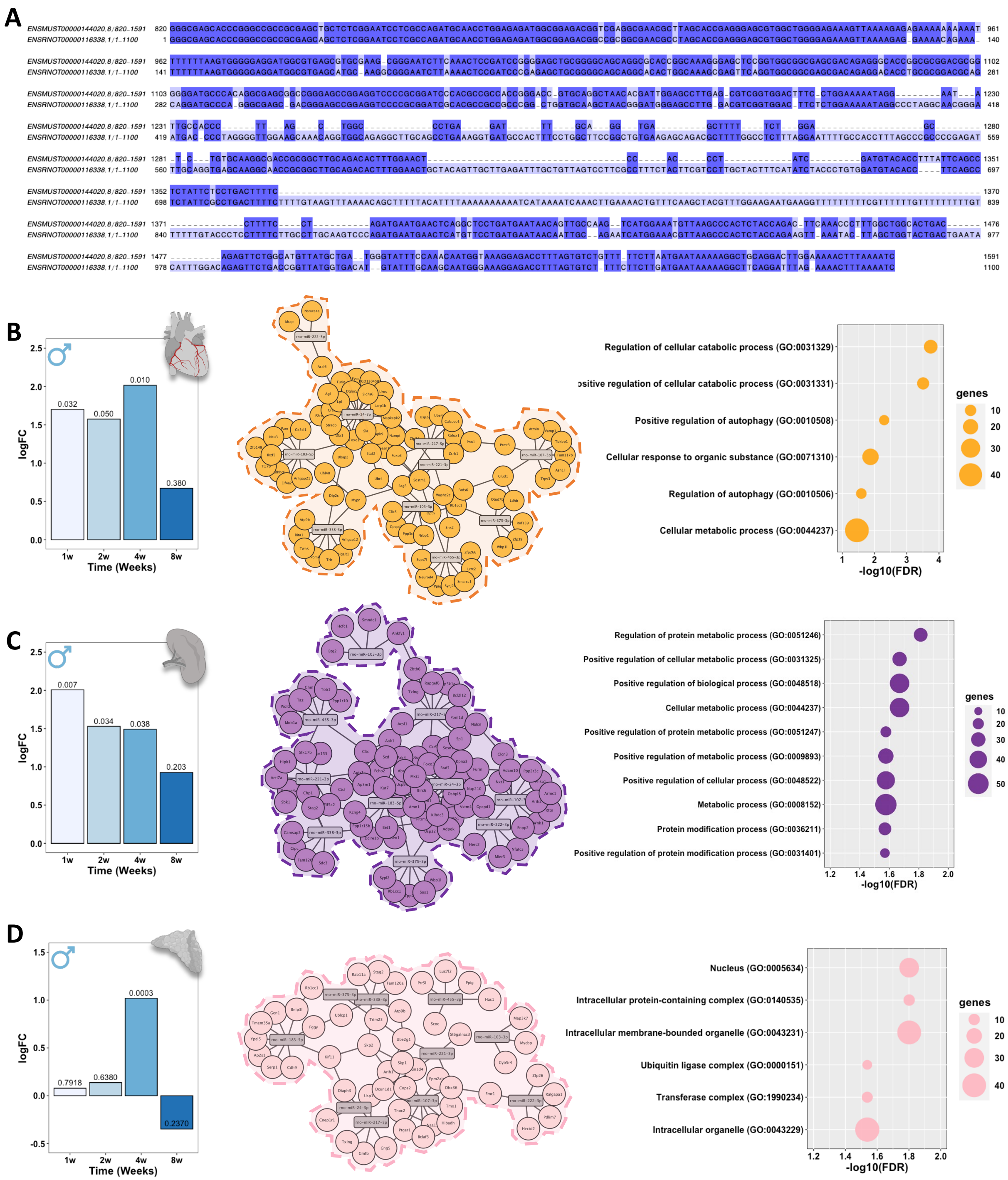


**Figure S7. Comparative analysis of the rat lncRNA ENSRNOG00000064728 and its ortholog ENSMUSG00000085837 in mice. (A)** RNA sequence conservation between rat and mouse transcripts. **(B)** Expression profile of lncRNA ENSRNOG00000064728 in male rat hearts (left panel), alongside its predicted miRNA-mRNA interaction network (center panel, in orange), and the associated Gene Ontology (GO) terms enriched from this network (right panel). **(C)** Expression profile of lncRNA ENSRNOG00000064728 in male rat spleens (left panel), with its predicted miRNA-mRNA interaction network (center panel, in purple), and the corresponding GO terms (right panel). **(D)** Expression profile of lncRNA ENSRNOG00000064728 in male rat adrenal glands (left panel), displaying its predicted miRNA-mRNA interaction network (center panel, in pink), and the relevant CC terms (right panel).

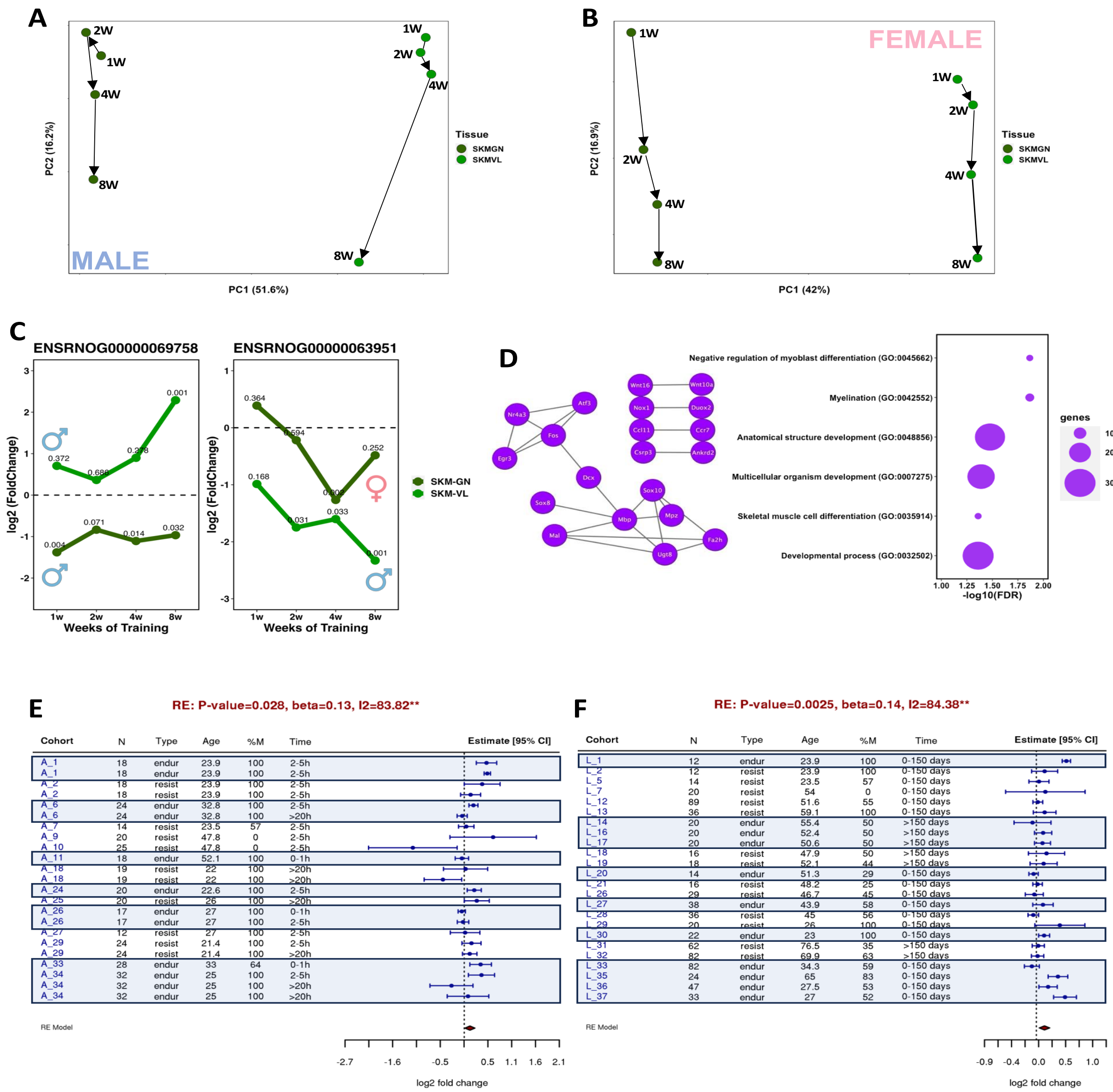

**Figure S8**. **Muscle-specific variation in lncRNA expression during an 8-week training regimen.**

**(A-B)** Principal Component Analysis (PCA) illustrating the distribution of male (**A**) and female (**B**) samples over the 8-week training period. **(C)** Temporal expression patterns of common lncRNAs (ENSRNOG00000069758 and ENSRNOG00000063951) in gastrocnemius (SKMGN) and vastus lateralis (SKMVL) tissues across training weeks. *p-values* indicate statistical significance at each stage of training. **(D)** Gene Ontology (GO) enrichment analysis of differentially expressed lncRNAs (DELncs) highlights key biological processes, including negative regulation of myoblast differentiation, myelination, anatomical structure development, and skeletal muscle cell differentiation. **(E-F)** APLN levels in skeletal muscle during acute (**E**) and chronic (**F**) endurance exercise in human subjects. Blue boxes represent values associated with endurance exercise.

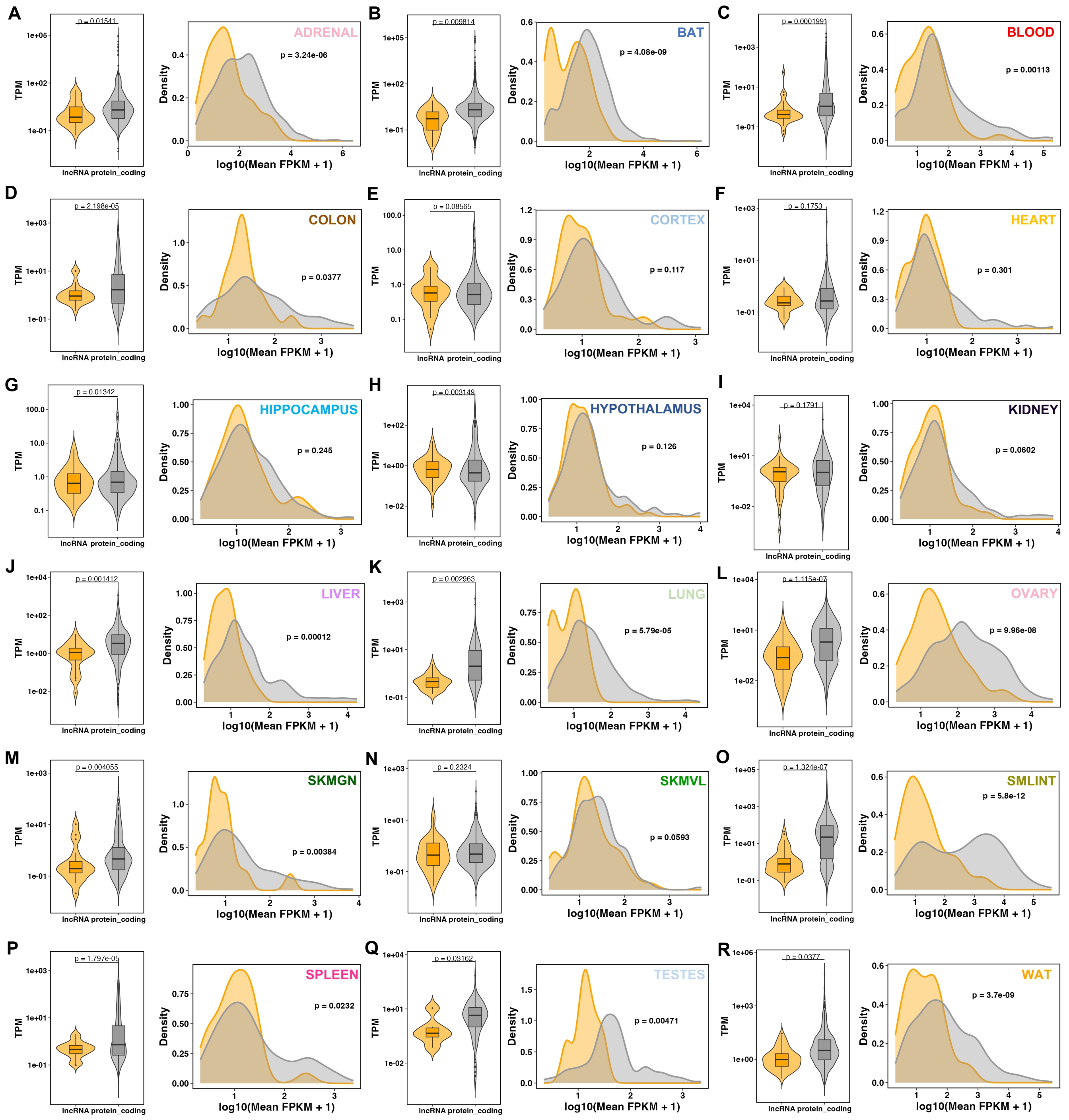


**Figure S9**. **Comparative expression analyses of lncRNAs and protein-coding genes across 18 rat tissues following exercise.**

Expression distributions were assessed using two complementary approaches: (left panel) boxplots of transcript per million (TPM), which allow direct comparison of relative expression levels between lncRNAs (orange) and protein-coding genes (grey); and (right panel) density plots of fragments per kilobase of transcript per million mapped reads (FPKM), which provide an overview of the overall distribution of transcript abundances. Statistical significance was evaluated using Welch's *t-test* (mean TPM) and the Kolmogorov–Smirnov test (FPKM distributions). *p-values* are indicated within each panel.

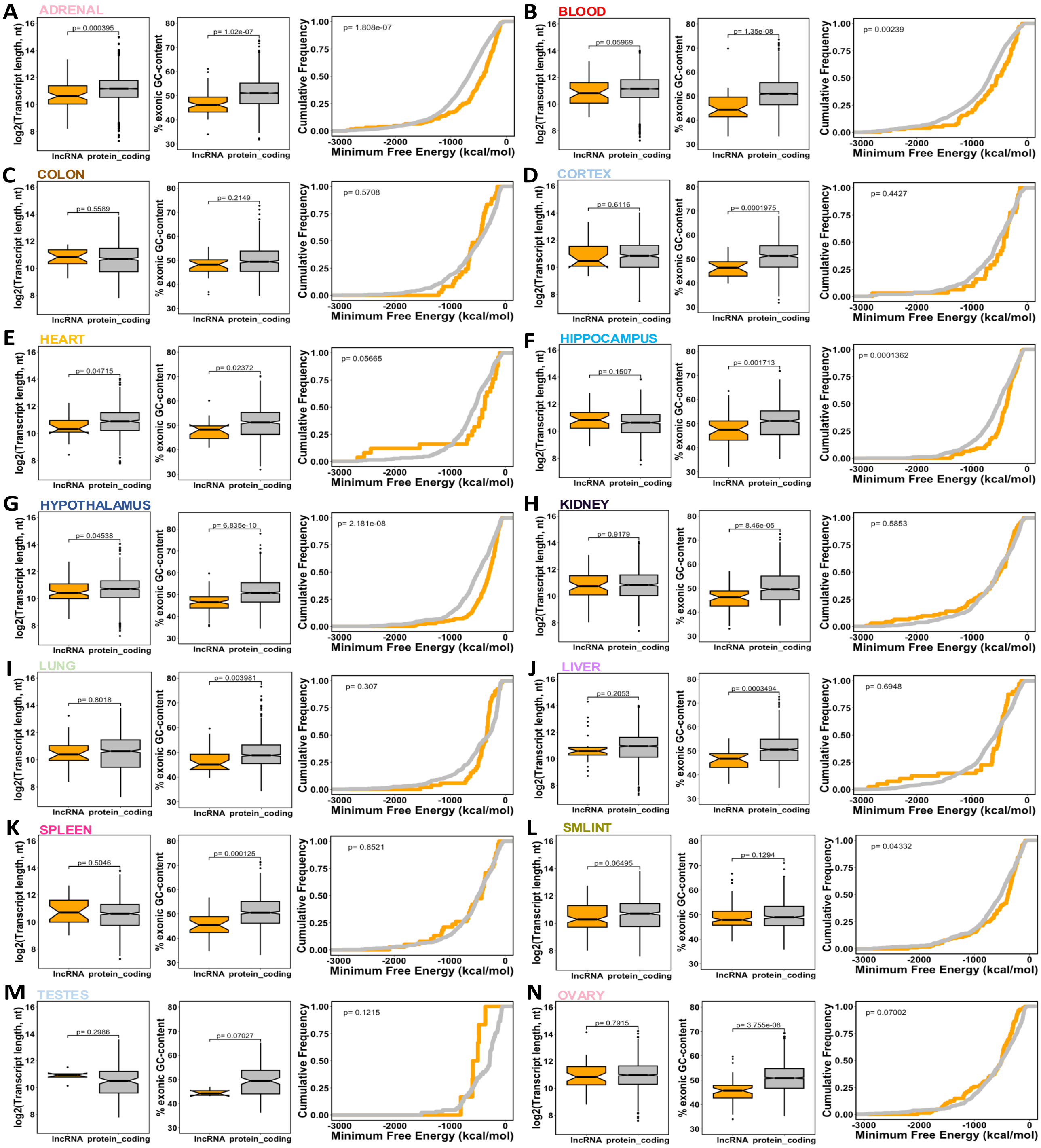


**Figure S10. Comparative analysis of lncRNA and protein-coding transcripts across various tissues.** (**A–N**) Tissue-specific comparisons of transcript length (left panel), GC content (center panel), and minimum free energy (MFE) (right panel) between lncRNAs (orange) and protein-coding transcripts (grey). Statistical significance was assessed using the Wilcoxon rank-sum test for transcript length and GC content, and the Kolmogorov–Smirnov test for MFE distributions. *p-values* are indicated in each panel.

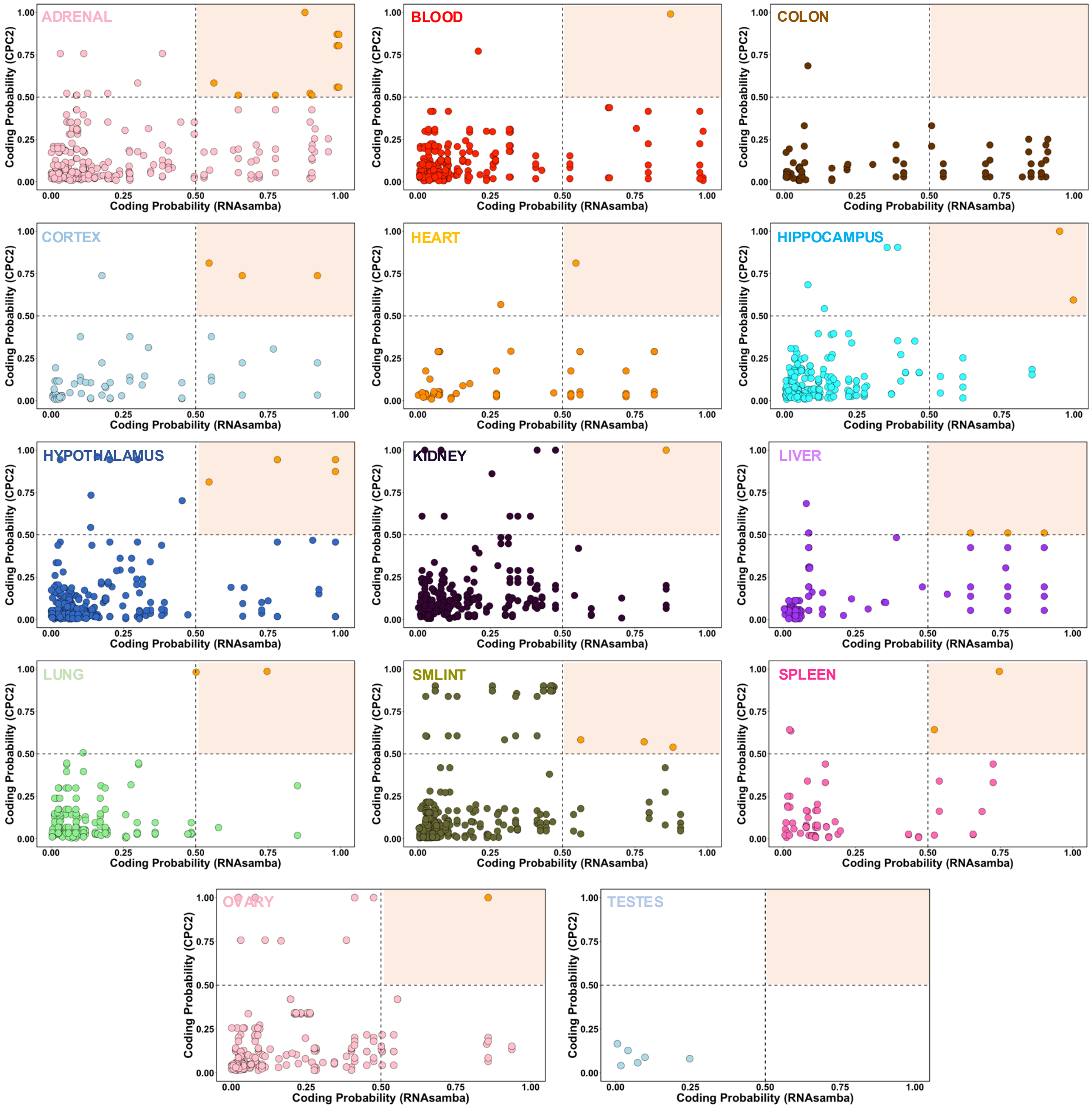


**Figure S11**. **Comparative analysis of coding probability among differentially expressed lncRNA isoforms across tissues.** Scatter plots illustrate the coding probability of lncRNA transcripts across various tissues, evaluated using RNAsamba and CPC2. Each panel represents a specific tissue, with RNAsamba-derived probabilities on the x-axis and CPC2-derived probabilities on the y-axis. Transcripts with scores < 0.5 in both analyses were classified as noncoding. These results indicate that the majority of analyzed transcripts are predicted to be noncoding.

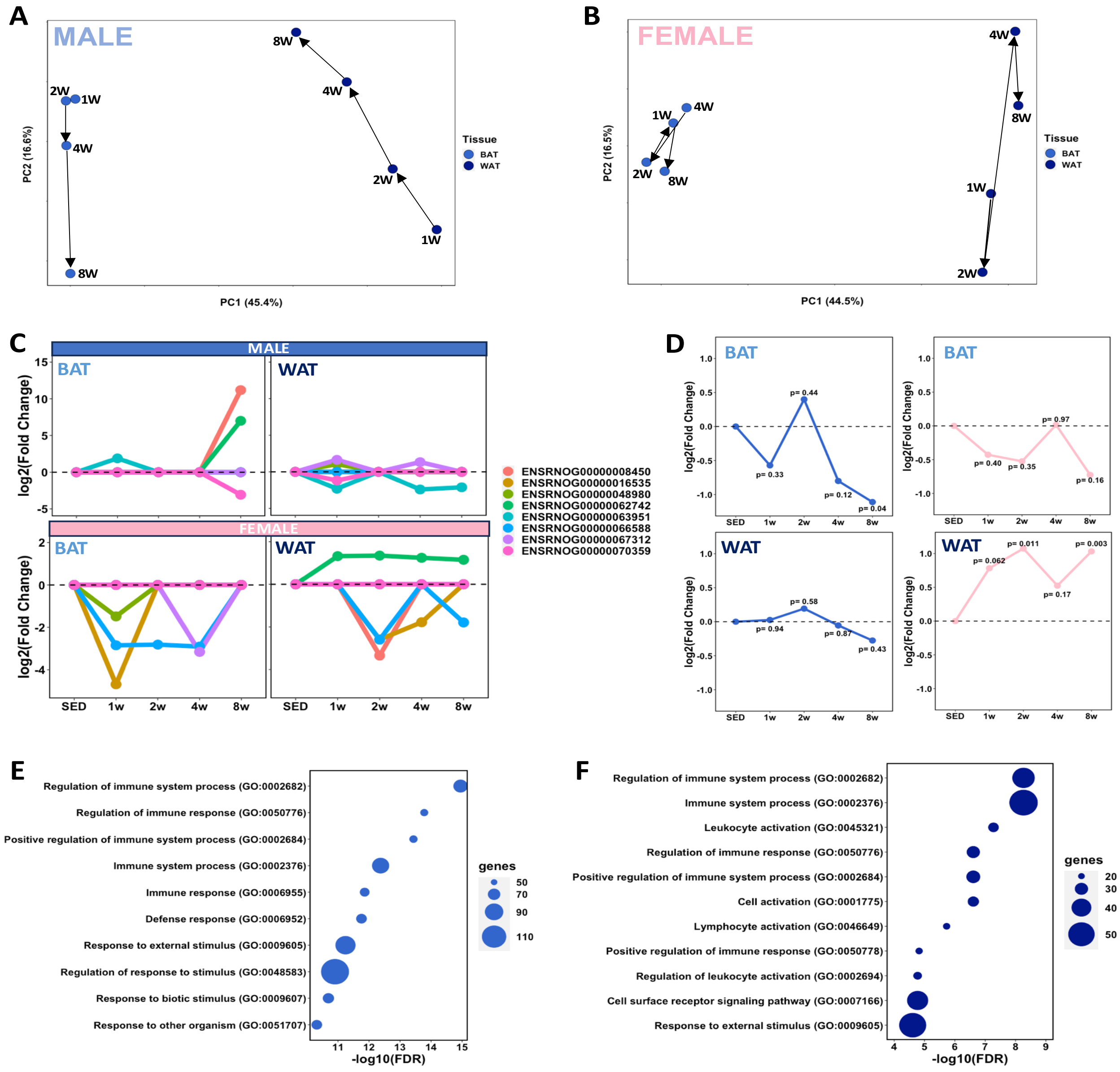


**Figure S12. Adipose tissue-specific variation in lncRNA expression during an 8-week training regimen. (A-B)** Principal Component Analysis (PCA) illustrating the distribution of male (**A**) and female (**B**) samples over the 8-week training period. **(C)** Temporal expression patterns of common differentially expressed lncRNAs (DELncs) in brown adipose tissue (BAT) and white adipose tissue (WAT) across training weeks. **(D)** Apelin expression dynamics in WAT of female rats, showing a significant increase following 8 weeks of endurance training. Blue and pink lines represent male and female rats, respectively. Statistical significance (*p-values*) is provided for each training stage. **(E-F)** Gene network correlation analysis between DELncs and differentially expressed protein-coding genes (DEGs) revealed gene ontology pathways in BAT (**E**) and WAT (**F**).

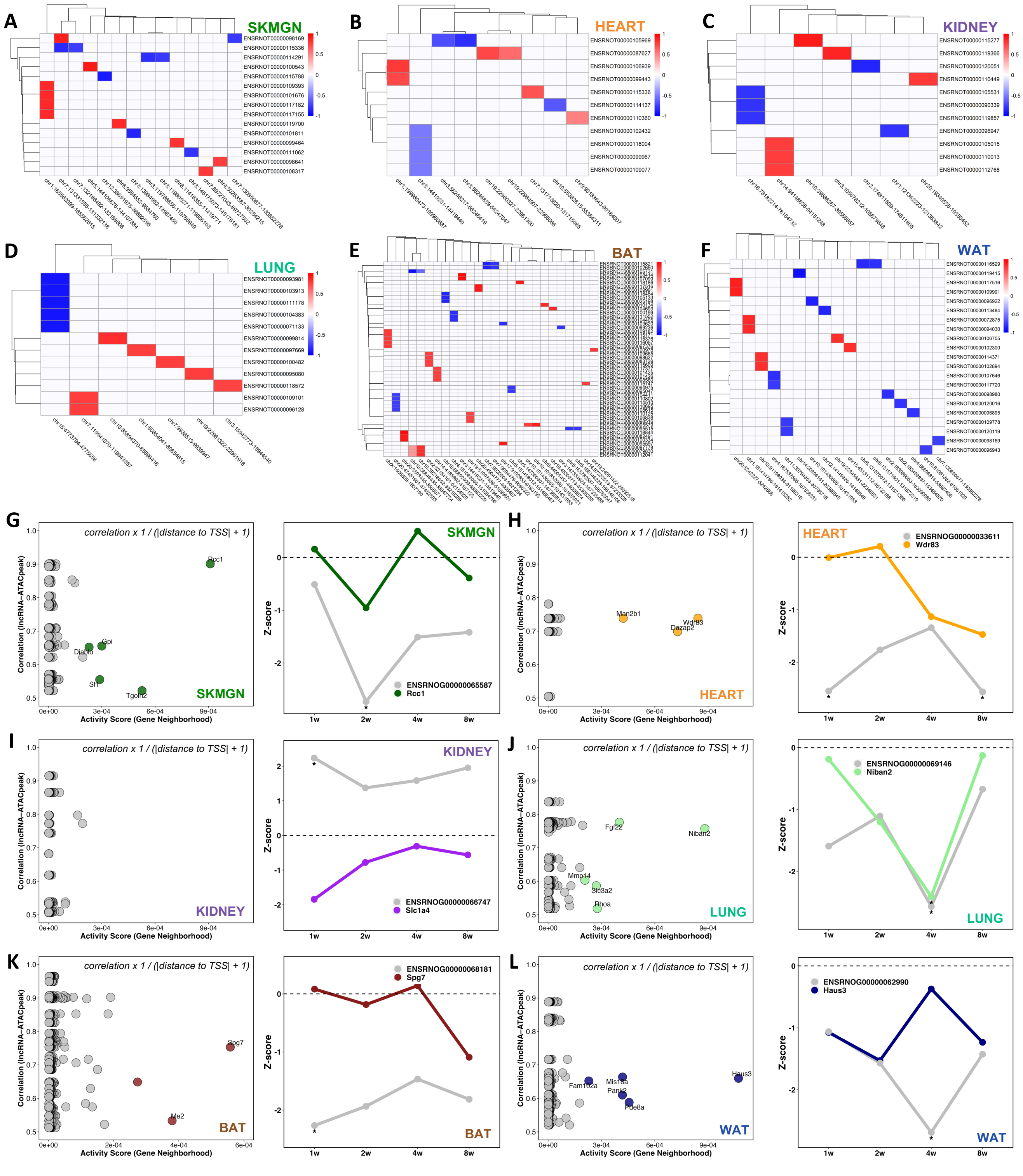


**Figure S13**. **Tissue-specific lncRNA–chromatin accessibility correlations and candidate cis-regulatory interactions. (A–F)** Heatmaps showing top significant correlations between differentially expressed lncRNAs and ATAC peaks in **(A)** SKMGN, **(B)** heart, **(C)** kidney, **(D)** lung, **(E)** BAT, and **(F)** WAT.

**(G–L)** Dot plots of activity scores (left) versus average correlation between lncRNAs and chromatin accessibility at neighboring gene promoters, highlighting candidate cis-regulatory lncRNAs in each tissue. Temporal expression profiles (Z-scores) of selected lncRNAs and their protein-coding gene pairs across training timepoints (right) are shown for **(G)** SKMGN, **(H)** heart, **(I)** kidney, **(J)** lung, **(K)** BAT, and **(L)** WAT.
